# Lis1 has two opposing modes of regulating cytoplasmic dynein

**DOI:** 10.1101/152645

**Authors:** Morgan E. DeSantis, Michael A. Cianfrocco, Zaw Min Htet, Phuoc Tien Tran, Samara L. Reck-Peterson, Andres E. Leschziner

## Abstract

Regulation is central to the functional versatility of cytoplasmic dynein, a motor involved in intracellular transport, cell division, and neurodevelopment. Previous work established that Lis1, a conserved and ubiquitous regulator of dynein, binds to its motor domain and induces a tight microtubule-binding state in dynein. The work we present here—a combination of biochemistry, single-molecule assays, cryo-electron microscopy and in vivo experiments—led to the surprising discovery that Lis1 has two opposing modes of regulating dynein, being capable of inducing both low and high affinity for the microtubule. We show that these opposing modes depend on the stoichiometry of Lis1 binding to dynein and that this stoichiometry is regulated by the nucleotide state of dynein’s AAA3 domain. We present data on the in vitro and in vivo consequences of abolishing the novel Lis1-induced weak microtubule-binding state in dynein and propose a new model for the regulation of dynein by Lis1.

## Introduction

Cytoplasmic dynein-1 (“dynein”) is the main microtubule-based motor that transports cellular cargos towards the minus-ends of microtubules, which are often located near the nucleus. In human and many other eukaryotic cells dynein distributes and organizes organelles, proteins, RNAs, and viruses, in addition to playing essential roles in cell division. Highlighting the fundamental nature of dynein, mutations in the dynein transport machinery cause a range of both neurodevelopmental and neurodegenerative diseases in humans (Franker and Hoogenraad, 2013).

For example, Type-1 lissencephaly is caused by mutations in one of dynein’s conserved regulators, Lis1 (*PAFAH1B1*). Lissencephaly is a rare but severe neurodevelopmental disease characterized by a smooth cerebral surface (Reiner et al., 1993). Lis1 was linked to the dynein pathway when mutations in the *Aspergillus nidulans* ortholog were shown to cause a nuclear distribution phenotype (Xiang et al., 1995), similar to mutations in the dynein motor in this filamentous fungus (Xiang et al., 1994). A number of mutations that cause defects in brain cortical development have since been mapped to the dynein motor-containing gene in humans (*DYNC1H1*) (Poirier et al., 2013). *S. cerevisiae* also has a Lis1 gene (Lee et al., 2003); however, in contrast to animals and filamentous fungi, dynein and Lis1 have a single, non-essential function in positioning the mitotic spindle in budding yeast, making it a powerful model system for dissecting dynein regulation by Lis1 (Huang et al., 2012; Roberts et al., 2014; Toropova et al., 2014).

There are conflicting models for the role that Lis1 plays during cargo transport. Lis1 has been implicated in many dynein-dependent functions (Cianfrocco et al., 2015; Kardon and Vale, 2009) such as (1) localizing and/or maintaining dynein at microtubule plus ends (Lee et al., 2003; Li et al., 2005; Sheeman et al., 2003; Splinter et al., 2012); (2) initiating dynein transport from microtubule plus ends (Egan et al., 2012; Lenz et al., 2006; Moughamian et al., 2013); and (3) enabling dynein to move against high loads (McKenney et al., 2010; Reddy et al., 2016; Yi et al., 2011). Other studies have looked at the effects of altering Lis1 expression levels on the flux and velocity of cargos transported by dynein. While most of these studies showed that deletion or depletion of Lis1 reduces transport (Dix et al., 2013; Klinman and Holzbaur, 2015; Moughamian et al., 2013; Pandey and Smith, 2011; Shao et al., 2013; Smith et al., 2000; Yi et al., 2011), at least one of them found that Lis1 depletion increases cargo transport (Vagnoni et al., 2016). It is not clear whether these apparent discrepancies reflect an incomplete understanding of how Lis1 regulates dynein or if Lis1’s activity is itself cargo-, organism-, or cell-specific.

At a mechanistic level, how does Lis1 regulate dynein? Because Lis1 binds directly to dynein’s motor domain (Huang et al., 2012; Toropova et al., 2014), it is essential to consider Lis1 regulation in the context of dynein’s mechanochemical cycle. The dynein motor is composed of two identical “heavy chains” of approximately 500 kDa each, and two copies each of 5 additional accessory subunits. Dynein’s heavy chain can be divided into two functional units: its N-terminal “tail”, which is required for dimerization, binding the accessory subunits, and cargo interactions (via additional proteins); and the “motor”, which is built around an AAA+ (**<u>A</u>**TPase **<u>A</u>**ssociated with diverse cellular **<u>A</u>**ctivities) ring containing 6 AAA domains (Figure 1A) (Carter, 2013; Cianfrocco et al., 2015). Dynein’s motility requires cycles of ATP hydrolysis at AAA1, which are coupled to cycles of microtubule binding and release at dynein’s microtubule binding domain (MTBD).

**Figure 1.**
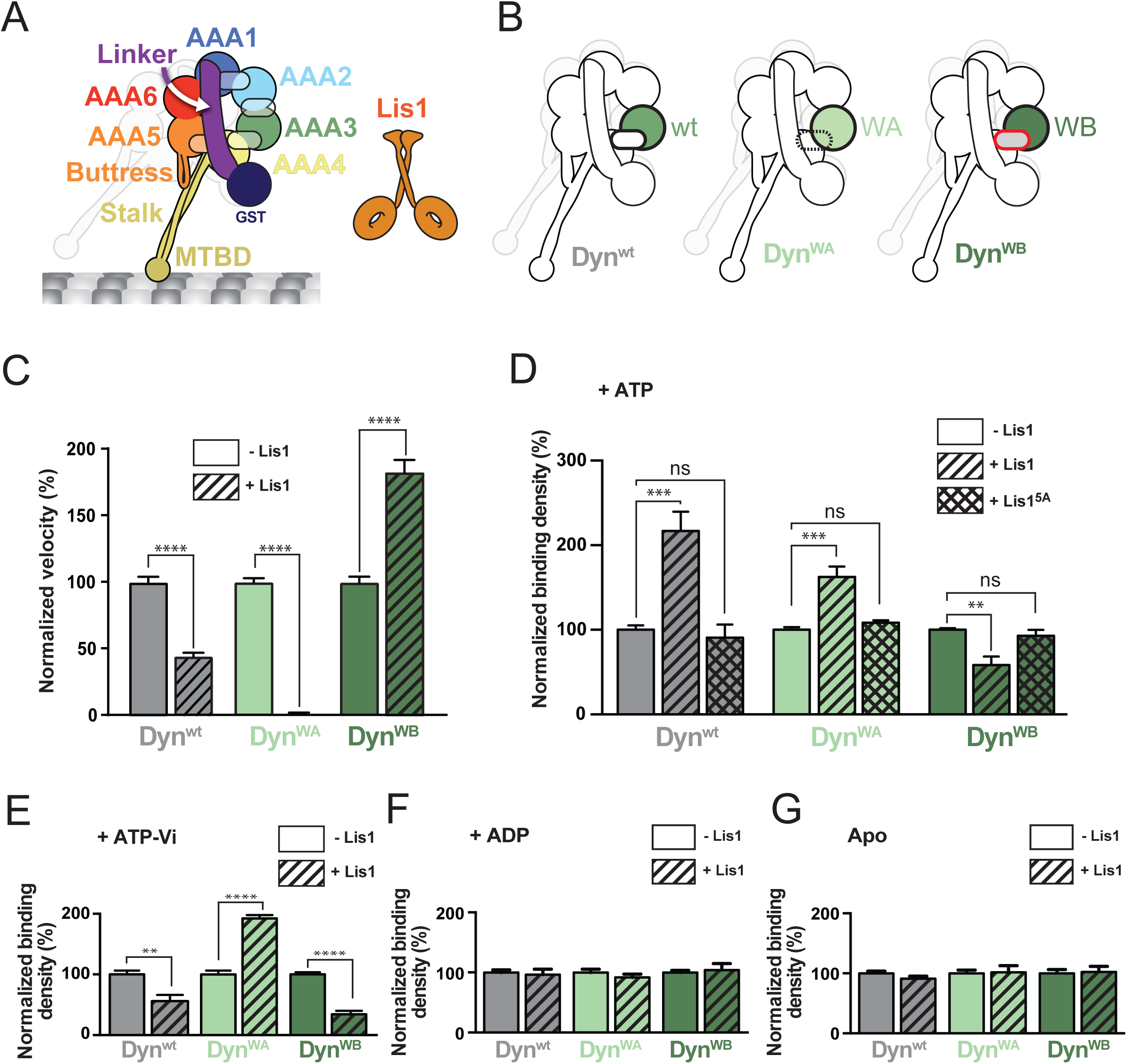
Lis1 has two modes of regulating dynein. (A) Schematic representation of the dynein construct used in this study. Domains are color-coded and labeled accordingly. The semi-transparent ovals represent the nucleotide-binding sites in AAA1-4. The second protomer, in the back, is faded out for clarity. The cartoon of a Lis1 dimer (orange) represents each ß-propeller domain as a ring. (B) AAA3 variants used in this study: Dyn^wt^ (green, wild type AAA3), Dyn^WA^ (light green, Walker A mutation in AAA3), and Dyn^WB^ (dark green, Walker B mutation in AAA3). (C) Normalized average velocities of dynein variants in the absence (solid bars) and presence (hatched bars) of 300 nM Lis1 (n>103 per data point). Velocity in the presence of Lis1 was normalized relative to the velocity in the absence of Lis1, which was set to 100%. (D-G) Normalized binding densities of dynein variants alone (solid bars), or in the presence of 300 nM Lis1 (hatched bars), or 300 nM Lis1^5A^ (cross-hatched bars). The nucleotide conditions in the imaging buffers were 1 mM ATP (D), 1 mM ATP-Vi (E), 1 mM ADP (F) and 2.5 units/mL apyrase (“Apo”) (G). For each variant, the binding density in the presence of Lis1 or Lis1^5A^ was normalized to dynein alone (see Methods) [n=12 (D), or n=8 (E-G) per data point]. Statistical significance was calculated using an unpaired t-test with Welch’s correction for both velocity (C) and binding density (D-G). P-values: ns, not significant; **, <0.01; ***, <0.001, ****, <0.0001. Data are shown as mean and standard error of mean.

Current evidence suggests that Lis1 acts on dynein by disrupting the coupling between these cycles of ATP hydrolysis and microtubule binding and release. Lis1 is a dimer of two ß-propellers (Tarricone et al., 2004), with a single ß-propeller binding to AAA4 on dynein’s ring (Huang et al., 2012) (Toropova et al., 2014). In vitro motility assays showed that Lis1 decreases dynein’s velocity (Huang et al., 2012; Yamada et al., 2008). Lis1 does this by causing dynein to remain tightly bound to microtubules, even in the presence of ATP (Huang et al., 2012; McKenney et al., 2010), which would normally result in microtubule release. This molecular model for the regulation of dynein by Lis1 is at odds with some of the proposed cellular functions of Lis1 discussed above. For example, depletion of Lis1 decreases the velocity of acidic organelles in mouse neurons (Pandey and Smith, 2011) and mRNAs in *Drosophila* embryos (Dix et al., 2013) even though a velocity increase would be expected when a factor that increases dynein’s affinity for microtubules is removed, as higher microtubule affinity directly correlates with lower velocity.

The regulation of dynein motility is complex, involving not only accessory proteins, such as Lis1, but also intra-motor control. In particular, the nucleotide state at AAA3 has been shown to regulate dynein’s motility by acting on its mechanochemical cycle (DeWitt et al., 2014; Nicholas et al., 2015). While stepping dynein contains ADP at AAA3, the presence of ATP or absence of nucleotide (“apo”) at this site results in dynein that is tightly bound to microtubules and exhibits a slowed velocity (DeWitt et al., 2014).This AAA3-mediated inhibitory effect occurs by keeping dynein bound to microtubules when ATP binds to AAA1, which normally triggers dynein’s release from microtubules (DeWitt et al., 2014).

We were struck by the similarities in the behavior of dynein in the presence of Lis1 or when its AAA3 is in an apo or ATP-bound state. The fact that Lis1’s binding site on AAA4 is close to AAA3’s ATP-binding site made us wonder if Lis1 regulates dynein by acting on those self-regulatory AAA3 states. We have used well-characterized mutants in *S. cerevisiae* dynein to control the nucleotide state of AAA3 (Cho et al., 2008; DeWitt et al., 2014; Nicholas et al., 2015) and a combination of single-molecule imaging, high-resolution cryo-electron microscopy (cryo-EM), *in vitro* reconstitutions, and *in vivo* assays to understand the relationship between Lis1 and the nucleotide state of dynein’s AAA3 domain. Our experiments led to the surprising discovery that Lis1 can regulate dynein in two distinct and opposing ways, depending on the nucleotide state of AAA3. We show that these two modes of regulation rely on different dynein:Lis1 ß-propeller stoichiometries: binding of a single ß-propeller to dynein with an AAA3 containing ADP or no nucleotide results in high affinity for microtubules, while binding of two ß-propellers to dynein with an AAA3 containing ATP results in a low affinity state. By mutating the newly discovered second binding site for Lis1 on dynein, we were able to eliminate the Lis1-induced low affinity state and probe the consequences of its removal. We propose a new model for the regulation of dynein by Lis1 that can now explain the multiple and conflicting cellular roles of Lis1.

## Results

### Lis1 has two modes of regulating dynein

Previous studies led us to hypothesize that the nucleotide state at AAA3 might play a role in the regulation of dynein by Lis1 (DeWitt et al., 2014; Nicholas et al., 2015; Toropova et al., 2014). To test this, we used a minimal *S. cerevisiae* dynein construct that is dimerized by GST (Dyn^wt^ hereafter) and has similar motile properties to full-length dynein (DeWitt et al., 2012; Gennerich et al., 2007; Reck-Peterson et al., 2006) (Figure 1A). also regulates Dyn^wt^ Importantly, Lis1 similarly to full-length dynein (Huang et al., 2012). In order to mimic different nucleotide states at AAA3 we made well-characterized mutations in it (Cho et al., 2008; DeWitt et al., 2014; Kon et al., 2004), which disrupt the conserved Walker A and Walker B motifs that distinguish the AAA+ family (Erzberger and Berger, 2006). The Walker A mutant (Dyn^WA^) impairs ATP binding, while the Walker B mutant (Dyn^WB^) allows ATP to bind, but prevents its hydrolysis (Figure 1B). We used these mutants throughout the work presented here to determine how Lis1 influences dynein activity when AAA3 is either in a nucleotide-free (Dyn^WA^) or ATP-bound (Dyn^WB^) state. Lis1 had a similar binding affinity for all three dynein variants (Figure S1A, Table S1).

We first asked if the nucleotide state at AAA3 altered Lis1’s effect on dynein’s velocity using single-molecule motility assays (Figure 1C, Table S1). In agreement with previous studies, Lis1 decreased the velocity of Dyn^wt^ (Huang et al., 2012; McKenney et al., 2010; Toropova et al., 2014; Yamada et al., 2008). While both Dyn^WA^ and Dyn^WB^ had slower velocities than Dyn^wt^ on their own (DeWitt et al., 2014) (Table S1), the relative effects of adding Lis1 to them were striking. Dyn^WA^ was hypersensitive to Lis1, with a velocity reduction of 99% relative to Dyn^WA^ alone (Figure 1C). Unexpectedly, the Lis1 effect was reversed with Dyn^WB^; the velocity of this mutant almost doubled in the presence of Lis1 (Figure 1C). These results suggest that Lis1 can regulate dynein’s velocity in opposite ways depending on the nucleotide state at AAA3.

Next, we asked if the nucleotide state at AAA3 altered Lis1’s effect on dynein’s affinity for microtubules. To measure microtubule binding affinities, we developed a single-molecule total internal reflection fluorescence (TIRF) microscopy-based assay to quantify the density of dyneins bound to microtubules (see Methods). Consistent with our previous work (Huang et al., 2012), we observed increased binding of Dyn^wt^ to microtubules in the presence of Lis1 (Figure 1D). In the absence of Lis1, both Dyn^WA^ and Dyn^WB^ had higher affinities for microtubules when compared to Dyn^wt^ (Figure S1B). In agreement with our velocity data (Figure 1C), the two AAA3 variants were also regulated by Lis1 in opposite ways: Lis1 increased the binding density of Dyn^WA^ by 62% and decreased the microtubule binding density of Dyn^WB^ by 42% (Figure 1D). Importantly, a Lis1 mutant we designed to abolish its interaction with dynein (Lis1^5A^) (Toropova et al., 2014) had no effect on any of the variants (Figure 1D), indicating that our observations are specific to the Lis1-dynein interaction. Thus, Lis1 can also regulate dynein’s microtubule binding in opposing manners depending on the nucleotide state at AAA3.

Given how unexpected Lis1’s effects on Dyn^WB^ were we wanted to repeat our single-molecule microtubule binding experiments with Dyn^wt^ trapped with AAA3 in a nucleotide state similar to that of Dyn^WB^ (ATP). To do this we used ATP plus Vanadate (ATP-Vi), which becomes ADP-Vi upon ATP hydrolysis (Burgess et al., 2003). In the presence of ATP-Vi, the AAA1 sites of Dyn^wt^, Dyn^WA^ and Dyn^WB^ are all expected to be in the ADP-Vi state (Schmidt et al., 2014). Their AAA3 sites, on the other hand, should reflect the nature of each variant, with Dyn^WA^’s AAA3 being empty, and Dyn^wt^ and Dyn^WB^ containing ADP-Vi and ATP, respectively. If the effect of Lis1 on microtubule binding reflects the nucleotide state of AAA3, then Dyn^wt^ and Dyn^WB^ might be expected to have a similar response to Lis1 in the presence of ATP-Vi. Indeed, under ATP-Vi conditions, Dyn^wt^ and Dyn^WB^ both showed decreased microtubule binding in the presence of Lis1 (Figure 1E). This was in striking contrast to the increased microtubule binding we saw for Dyn^wt^ with ATP (Figure 1D), where AAA3 is expected to be mainly in an ADP state (DeWitt et al., 2014). The affinity of Dyn^WA^, which cannot bind nucleotide at AAA3, still increased in the presence of ATP-Vi and Lis1 (Figure 1E). Performing these same experiments in the presence of ADP or in the absence of nucleotide abolished all differences among the three variants (Figure 1F, G), suggesting that dynein’s AAA1 must be in an ATP or ADP-Pi state for Lis1 regulation to be apparent.

Together, these results reveal that in addition to the previously characterized ability of Lis1 to promote tight binding to microtubules by dynein (Huang et al., 2012; McKenney et al., 2010; Toropova et al., 2014), Lis1 can also promote weak binding of dynein to microtubules. Our data suggest that the nucleotide state of AAA3 determines which of these opposing modes of regulation Lis1 uses: an AAA3 in an ADP or apo state results in Lis1 inducing strong binding to microtubules, while a AAA3 in an ATP or ADP-Pi state results in Lis1 inducing weak binding to microtubules. We next sought to understand the mechanistic basis for this surprising dual role of Lis1 by determining the structures of dynein:Lis1 complexes using different AAA3 variants.

### Structural basis for the tight microtubule binding state of dynein induced by Lis1

We begin this section with a brief description of some aspects of dynein’s structure that are relevant to the results presented below. More detailed reviews of dynein’s structure can be found elsewhere (Carter, 2013; Cianfrocco et al., 2015; Gleave et al., 2014).

As in other AAA+ ATPases, the AAA domains in dynein are comprised of “small” and “large” subdomains (Figure 2A) (Erzberger and Berger, 2006). We refer to these subdomains as AAA“X” S and AAA“X”L, with “X” denoting the number of the AAA domain to which they belong. The small and large subdomains are arranged in two planes. Dynein’s linker forms a third plane, resulting in an overall order of linker, large subdomains, small subdomains (Figure 2A). ATP binding and hydrolysis take place at the boundary between adjacent AAA modules. The Walker A and B motifs that enable binding and hydrolysis are located in the large subdomain, while the following large subdomain contains an arginine residue (the “arginine finger”) that is required for ATP hydrolysis (Gleave et al., 2014). Conformational changes in dynein’s ring alter the nucleotide-binding sites and regulate the affinity of the motor for its track, the microtubule (Carter, 2013; Cianfrocco et al., 2015). High affinity states of dynein are associated with an opening of the ring while low affinity states are associated with its compaction (Kon et al., 2012; Schmidt et al., 2014; 2012). The coupling between ring conformations and changes in affinity at the microtubule-binding domain is mediated by dynein’s stalk, a long antiparallel coiled coil, and the buttress, a short antiparallel coiled coil that comes out of AAA5 and interacts with one of the helices in the stalk (Figure 2A). The position of the buttress and the register of the stalk’s coiled coil change throughout dynein’s mechanochemical cycle, leading to different conformations and affinities of the microtubule-binding domain (Carter et al., 2008; Gibbons, 2005; Kon et al., 2009; Redwine et al., 2012; Schmidt et al., 2014).

**Figure 2.**
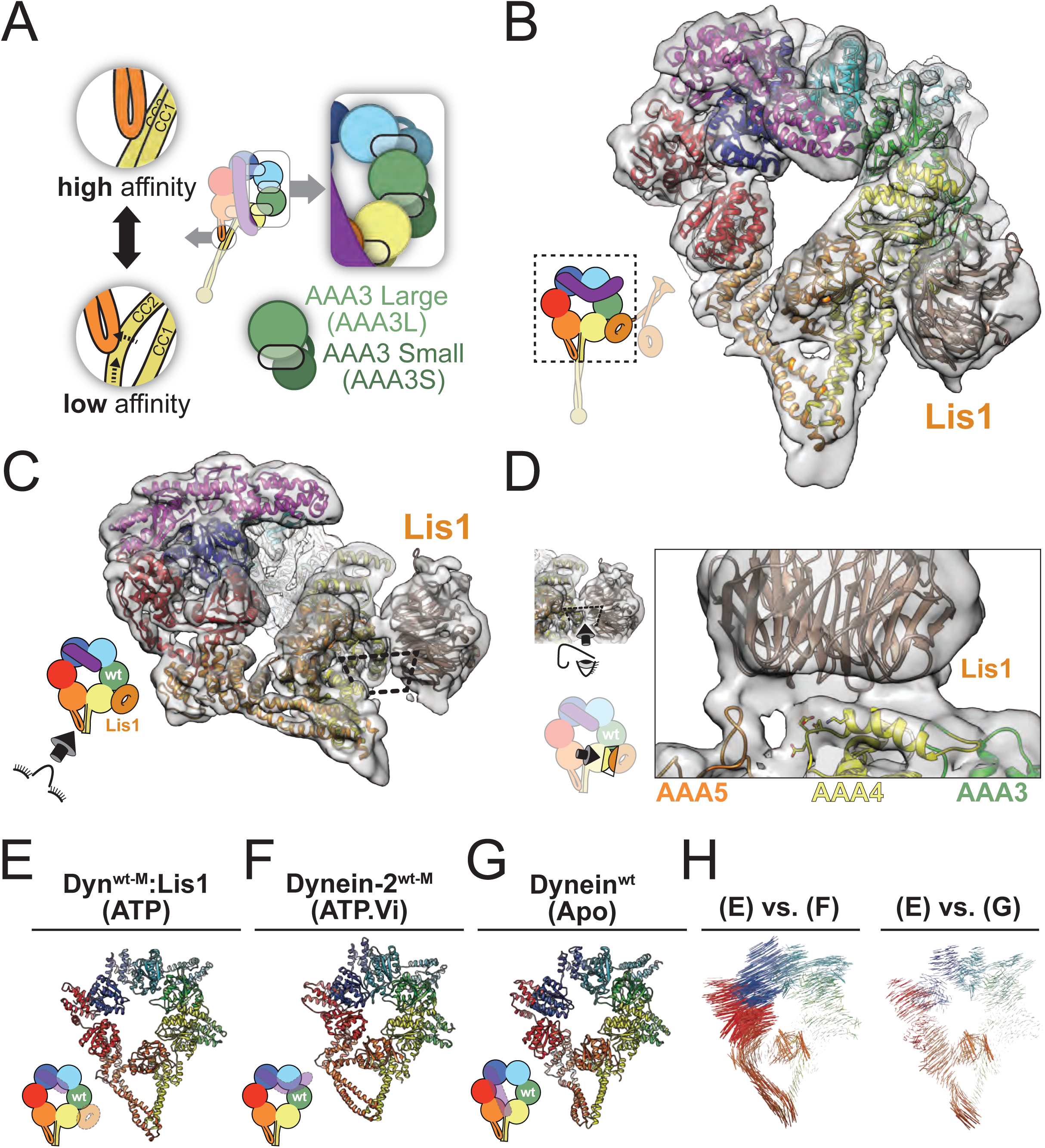
Structural basis for the tight microtubule binding state of dynein induced by Lis1. (A) This panel highlights in more detail some structural features of the schematic shown in Figure 1A. The inset on the right shows the architecture of the AAA ring, with each AAA domain composed of both large (AAAL) and small (AAAS) subdomains (as exemplified for AAA3). The large and small subdomains are arranged in two separate planes, with lighter and darker shades used in the inset to represent their positions closer to or further away from the viewer, respectively. The semitransparent ovals represent the nucleotide-binding sites. The insets on the left illustrate how the buttress couples the conformation of dynein’s ring to dynein’s affinity for microtubules by changing the register between the two helices (CC1 and CC2) in the stalk’s coiled coil. (B) Cryo-EM structure of the Dyn^wt-M^:Lis1 complex, solved in the presence of ATP. The overall 7.7Å-resolution cryo-EM map was filtered using local resolution values and is shown as a semi-transparent surface, with the atomic model generated with Rosetta shown as a ribbon diagram. The schematic representation (bottom left) indicates the portion of the dynein monomer that was observed in our cryo-EM map. (C) In this panel the structure is shown viewed from the with the stalk, with the Dyn^wt-M^-Lis1 interface indicated by the dashed rectangle. (D) Close-up view of the Dyn^wt-M^-Lis1 interface, seen in a direction perpendicular to the dashed rectangle. Side chains shown on AAA4 are residues that prevent Lis1 binding when mutated (KDEE) (Huang et al., 2012). The AAA domains that contribute motifs to the interface are labeled. (E-G) Ring conformations of the Dyn^wt-M^:Lis1 structure (E); the low-affinity, wild-type human dynein-2 solved in the presence of ATP-Vi (PDB: 4RH7) (F); and the high-affinity, wild-type *S. cerevisiae* dynein solved in the absence of nucleotide (PDB: 4AKI) (G). We removed the linker and, when present, Lis1 from these representations in order to make the ring conformations easier to visualize. (H) Maps of pairwise alpha carbon interatomic distances between the Dyn^wt-M^:Lis1 structure and the low-affinity, wild-type human dynein-2 (left), and high-affinity, wild-type *S. cerevisiae* dynein (right). We calculated the pairwise alpha carbon interatomic distances shown here after aligning the structures using their AAA4L domains. The length and thickness of the vectors are proportional to the calculated interatomic distances.

To understand the structural basis of Lis1’s ability to increase dynein’s affinity for microtubules, we used single particle cryo-EM to determine the structure of Dynein:Lis1 in the presence of ATP, a state that has a high affinity for microtubules. For these studies, we used a monomeric dynein construct lacking GST (Dyn^wt-M^ hereafter), which is otherwise identical to the dimeric dynein constructs introduced above. Using cryo-EM, we determined a map of Dyn^wt-M^ bound to Lis1 in the presence of ATP at an overall resolution of 7.7Å and built an atomic model into the density using Rosetta (Figure 2B, Figure S2, and Table S2) (DiMaio et al., 2015; Wang et al., 2016).

This map, at much higher resolution than our previous 21Å structure of the Dyn^wt-M^:Lis1 complex (Toropova et al., 2014), provided a detailed view of the interface between dynein and Lis1 and revealed the conformation of dynein’s ring. As before, the structure showed a single Lis1 ß-propeller bound to Dyn^wt-M^, even though Lis1 was present as a dimer in our sample. The EM density suggested that Lis1 interacts not only with an alpha helix in AAA4, as we had previously described (Toropova et al., 2014), but also with a loop from AAA5, and possibly another from AAA3 (Figure 2C, D), although the resolution of the map does not establish this last interaction unambiguously.

Next we examined how Lis1 affected the conformation of dynein’s ring by comparing the AAA ring geometry of our Dyn^wt-M^:Lis1 structure (Figure 2E) to existing structures of dynein (Schmidt et al., 2012; 2014) (Figure 2F, G). Since we obtained our structure in the presence of ATP, where most AAA3s should contain ADP (DeWitt et al., 2014), one might have expected dynein’s ring to adopt a closed conformation reminiscent of that observed in the crystal structure of human dynein-2, which was solved in the presence of ATP-Vi (Figure 2F). However, in the presence of Lis1 and ATP, dynein adopts an open ring conformation, more similar to that observed for the nucleotide-free, tight-binding state of the motor (Schmidt et al., 2012) (Figure 2G). An analysis of interatomic distances between alpha carbons in our structure of Dyn^wt-M^:Lis1 and those in the structures of human dynein-2 (ATP-Vi) (Schmidt et al., 2014) and nucleotide-free *S. cerevisiae* dynein (Schmidt et al., 2012) illustrated the opening of the dynein ring upon Lis1 binding and highlighted the similarities between the Dyn^wt-M^:Lis1 and nucleotide-free dynein structures (Figure 2H).

Thus, our structural data show that a single Lis1 ß-propeller binding to dynein in the presence of ATP, which normally releases dynein from microtubules, results in a conformation associated with dynein having a high affinity for microtubules.

### Two Lis1 ß-propellers are bound to dynein in the weak microtubule binding state

After determining the structure of Dyn^wt-M^:Lis1, we moved on to the most puzzling aspect of our data: why does introducing a Walker B mutation in AAA3 (mimicking an ATP-bound state) lead to such a dramatic difference in how Lis1 regulates dynein? For this, we solved a 10.5Å-resolution cryo-EM structure of the Dyn^WB-M^:Lis1 complex in the presence of ATP-Vi. As in the Dyn^wt-M^:Lis1 structure, Dyn^WB-M^ is a monomeric construct lacking GST and Lis1 is a dimer. We again used Rosetta to build an atomic model of the complex (Figure 3A, Figure S3, and Table S2) (DiMaio et al., 2015; Wang et al., 2016).

**Figure 3.**
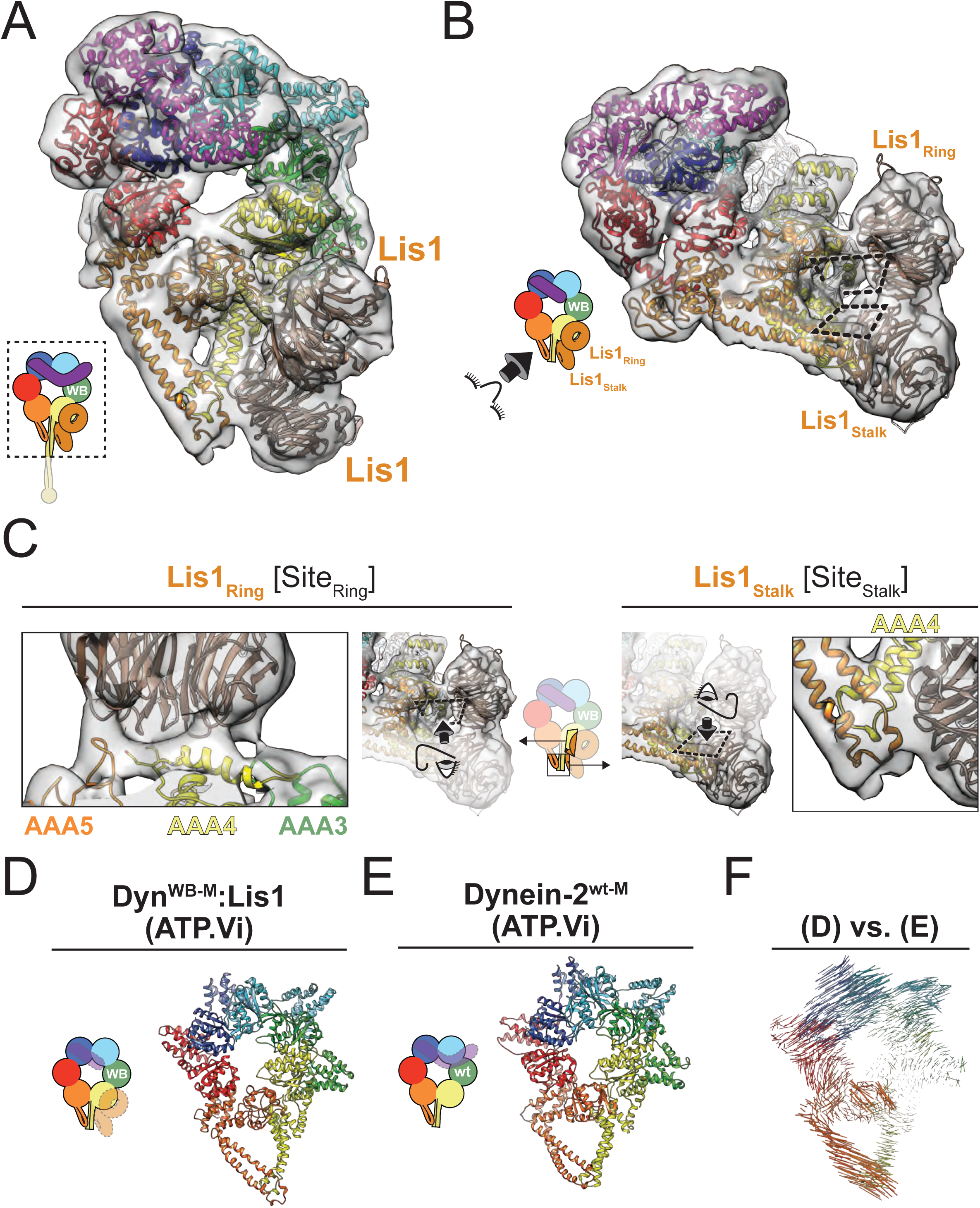
Two Lis1 β-propellers are bound to dynein in the weak microtubule binding state. (A) Cryo-EM structure of the Dyn^WB-M^:Lis1 complex, solved in the presence of ATP-Vi. The 10.5Å-resolution cryo-EM map is shown as a semi-transparent surface, with the atomic model generated with Rosetta shown as a ribbon diagram. The schematic representation (bottom left) indicates the portion of the dynein monomer that was observed in our cryo-EM map. (B) In this panel the structure is shown viewed from the stalk, with the two Dyn^WB-M^-Lis1 interfaces indicated by the dashed rectangles. (C) Close-up views of the two Dyn^WB-M^-Lis1 interfaces seen in the structure, viewed perpendicular to the dashed rectangles. The Lis1-binding site located on dynein’s ring (Site_Ring_) is shown on the left and the Lis1-binding site located on the stalk (Site_Stalk_) is shown on the right. The AAA domains that contribute motifs to the interfaces are labeled. (D-E) Ring conformations of the Dyn^WB-M^:Lis1 structure (D) and the low-affinity, wild-type human dynein-2 solved in the presence of ATP-Vi (PDB: 4RH7) (E). We removed the linker and, when present, Lis1 from these representations in order to make the ring conformations easier to visualize. (F) Map of pairwise alpha carbon interatomic distances between the Dyn^WB-M^:Lis1 structure and the low-affinity, wild-type human dynein-2. We calculated pairwise alpha carbon interatomic distances after aligning the structures using their AAA3L domains. The length and thickness of the vectors are proportional to the calculated interatomic distances.

Surprisingly, the cryo-EM map of Dyn^WB-M^:Lis1 showed not one, but two Lis1 11- propellers bound to dynein (Figures 3A, B). One of the 11-propellers binds to the previously identified site on the ring (AAA4) (“Site_Ring_”), while the other binds to dynein’s stalk, specifically coiled-coil 1 (CC1) (“Site_Stalk_”) (Figure 3A-C). Each Lis1 uses a different surface to interact with dynein, with the Lis1 at Site_Ring_ interacting with dynein across the narrower face of its ß-propeller, and Lis1 at Site_Stalk_ using the edge of its ß-propeller (Figure 3C). The density encompassing the two Lis1 ß-propellers is contiguous, suggesting they may interact with each other (Figure 3B), although our current resolution does not allow us to determine their rotational orientations and therefore the nature of this interface. Analysis of the stoichiometry of our Dyn^wt-M^:Lis1 complex using size-exclusion chromatography suggests that both ß-propellers in the structure belong to the same Lis1 dimer (Figure S4), as opposed to two separate Lis1 dimers.

The conformation of dynein’s ring in the Dyn^WB-M^:Lis1 structure is consistent with the weak microtubule binding of the complex. In this structure, dynein’s ring is in a closed conformation, similar to that seen in the crystal structure of the low microtubule affinity state of human dynein-2, solved in the presence of ATP-Vi (Figure 3D, E) (Schmidt et al., 2014). An analysis of interatomic distances between alpha carbons in our Dyn^WB-M^:Lis1 structure and those in the structure of human dynein-2 (ATP-Vi) illustrates the similarity between the conformations of their rings (Figure 3F); in contrast, a comparison between Dyn^WT-M^:Lis1 and human dynein-2 (ATP-Vi) revealed larger differences (Figure 2H).

Our results thus far show that the nucleotide state of AAA3 governs how Lis1 regulates dynein’s affinity for microtubules. Our biochemical data (Figure 1) indicate that the absence of nucleotide or the presence of ADP at AAA3 leads to a Lis1-induced tight binding state, while the presence of ATP results in a Lis1-induced weak binding state. Structurally, the nucleotide state of AAA3 is associated with different stoichiometries between dynein and Lis1 ß-propellers, with a single ß-propeller bound to dynein’s ring when AAA3 is empty (Toropova et al., 2014) or is occupied by ADP (Figure 2), and surprisingly two ß-propellers when AAA3 contains ATP (Figure 3).

### Lis1’s opposite modes of dynein regulation are associated with rigid body motion conformational changes in dynein’s ring

We wanted to understand how structural changes in dynein’s ring relate to the two modes of Lis1 regulation, and how the second Lis1 binding site at Site_Stalk_ in Dyn^WBM^:Lis1 stabilizes a low affinity state in dynein. In order to make this comparison, we calculated inter-atomic distances between Dyn^wt-M^:Lis1 and Dyn^WB-M^:Lis1 (Figure 4A). Unexpectedly, the resulting distance measurements showed that all of the differences were captured by the rigid motion of elements on one half of dynein’s ring—comprising the stalk, AAA5, AAA6 and part of AAA1—relative to the rest (Figure 4A, B).

**Figure 4.**
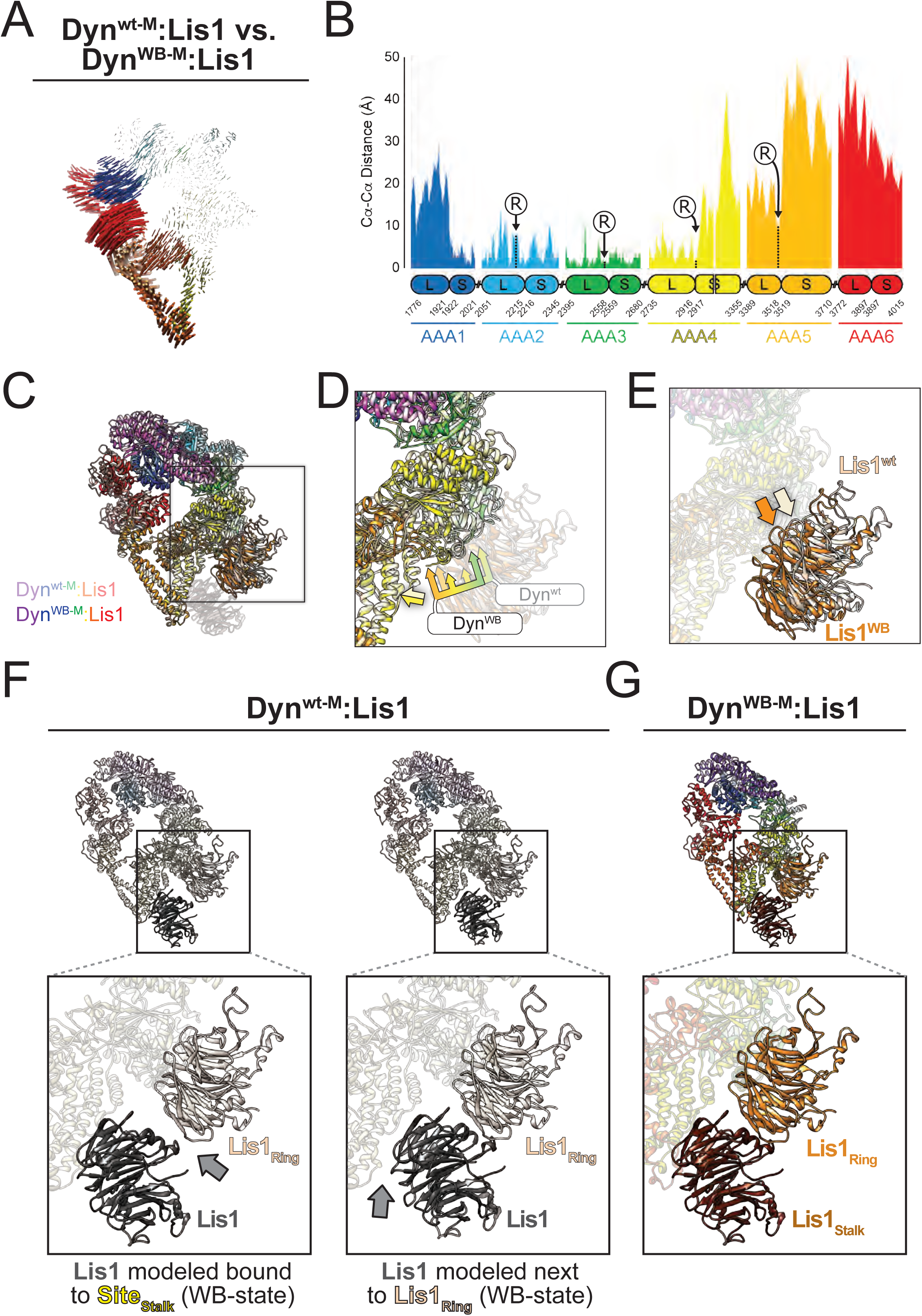
Lis1’s opposite modes of dynein regulation are associated with rigid body motion conformational changes in dynein’s ring. (A) Map of pairwise alpha carbon interatomic distances between the Dyn^wt-M^:Lis1 and Dyn^WB-M^:Lis1 structures. We removed the linker and Lis1 in order to make the ring conformations easier to visualize. We calculated pairwise alpha carbon interatomic distances after aligning the structures using their AAA3L domains. The length and thickness of the vectors are proportional to the calculated interatomic distances. (B) 1D plot of the interatomic distances shown in panel (A). The large and small subdomains in each AAA domain are indicated below the plot, along with the amino acid numbers at their boundaries. Distances are colored by AAA domain. The positions of the arginine fingers of domains AAA2-5, which act on domains AAA1-4, are labeled (“R”). (C) Superposition of the Dyn^wt-M^:Lis1 and Dyn^WBM^:Lis1 structures, aligned using their AAA4S domains (labeled). The Dyn^wt-M^:Lis1 structure is shown in lighter colors and the Lis1 bound to Site_Stalk_ in Dyn^WB-M^:Lis1 was faded for clarity. The square highlights Site_Ring_, and is the area represented in panels (D) and (E). (D) Close-up of the Site_Ring_ elements in dynein, with Lis1 faded for clarity. The bi-color arrows indicate good alignment for both AAA3S (green) and the base of the stalks (yellow colors). The tri-color multiheaded arrows point to the AAA3, AAA4 and AAA5 elements in Site_Ring_ in both Dyn^wt-M^:Lis1 (light colors) and Dyn^WB-M^:Lis1 (full colors). (E) The positions of the Site_Ring_-bound Lis1 in the Dyn^wt-M^:Lis1 and Dyn^WB-M^:Lis1 structures. This is the same view as in (D) but this time with dynein faded for clarity. The light and dark orange arrows point to the equivalent positions in Lis1 in the Dyn^wt-M^:Lis1 and Dyn^WB-M^:Lis1 structures, respectively. (F) Modeling of a second Lis1 into the Dyn^wt-M^:Lis1 structure. Lis1 was modeled either interacting with Site_Stalk_ (left) or with the Lis1 bound at Site_Ring_. The grey arrows point to the resulting gaps between the modeled Lis1 and Lis1 bound to Site_Ring_ (left) or between the modeled Lis1 and Site_Stalk_ (right). (G) For comparison, we show the same view of the experimentally observed Dyn^WB-M^:Lis1 structure.

The conformational change illustrated in Figure 4A results in the two Lis1 binding sites (Site_Ring_ and Site_Stalk_) being closer to each other in Dyn^WB-M^:Lis1 relative to Dyn^wt-M^:Lis1 (Figure 4C-E). The shorter distance between the two Lis1 binding sites in Dyn^WB-M^:Lis1 means that the interactions among dynein and the two Lis1’s cannot be satisfied in the Dyn^wt-M^:Lis1 structure, where we only observed a single Lis1 ß-propeller bound. This can be observed by modeling a second Lis1 ß-propeller into the Dyn^wt-M^:Lis1 structure, either bound to Site_Stalk_ (Figure 4F, left), or interacting with the Lis1 at Site_Ring_ (Figure 4F, right). Both models result in gaps of ~15 Å that do not exist in the Dyn^WB-M^:Lis1 structure (Figure 4G). We discuss below how these differences could play a role in the formation of the Dyn^WB-M^:Lis1 complex.

### The second Lis1 binding site is conserved in organisms that have a Lis1 orthologue in their genomes

The resolution of our cryo-EM map of the Dyn^WB-M^:Lis1 complex is not sufficient to unambiguously dock Lis1 and determine which residues interact with Site_Stalk_ in dynein. On the other hand, the segment of CC1 in dynein’s stalk where Site_Stalk_ must be located is short, limiting the number of candidate residues in dynein for this binding site (Figure 3C). When we performed an alignment of dynein sequences corresponding to this region of the stalk, we identified three residues—corresponding to E3012, Q3014 and N3018 in the *S. cerevisiae* sequence (Figure 5A, B)—that are conserved from yeast to humans (Figure 5A). However, the alignment also revealed a subset of fission yeasts where this conservation was not present (Figure 5A). Interestingly, and supporting the idea that these residues are central features of Site_Stalk_ in dynein, none of these fission yeasts contain a Lis1 orthologue in their genome, while the organisms with a conserved EQN triad contain Lis1 orthologues (Figure 5A).

**Figure 5.**
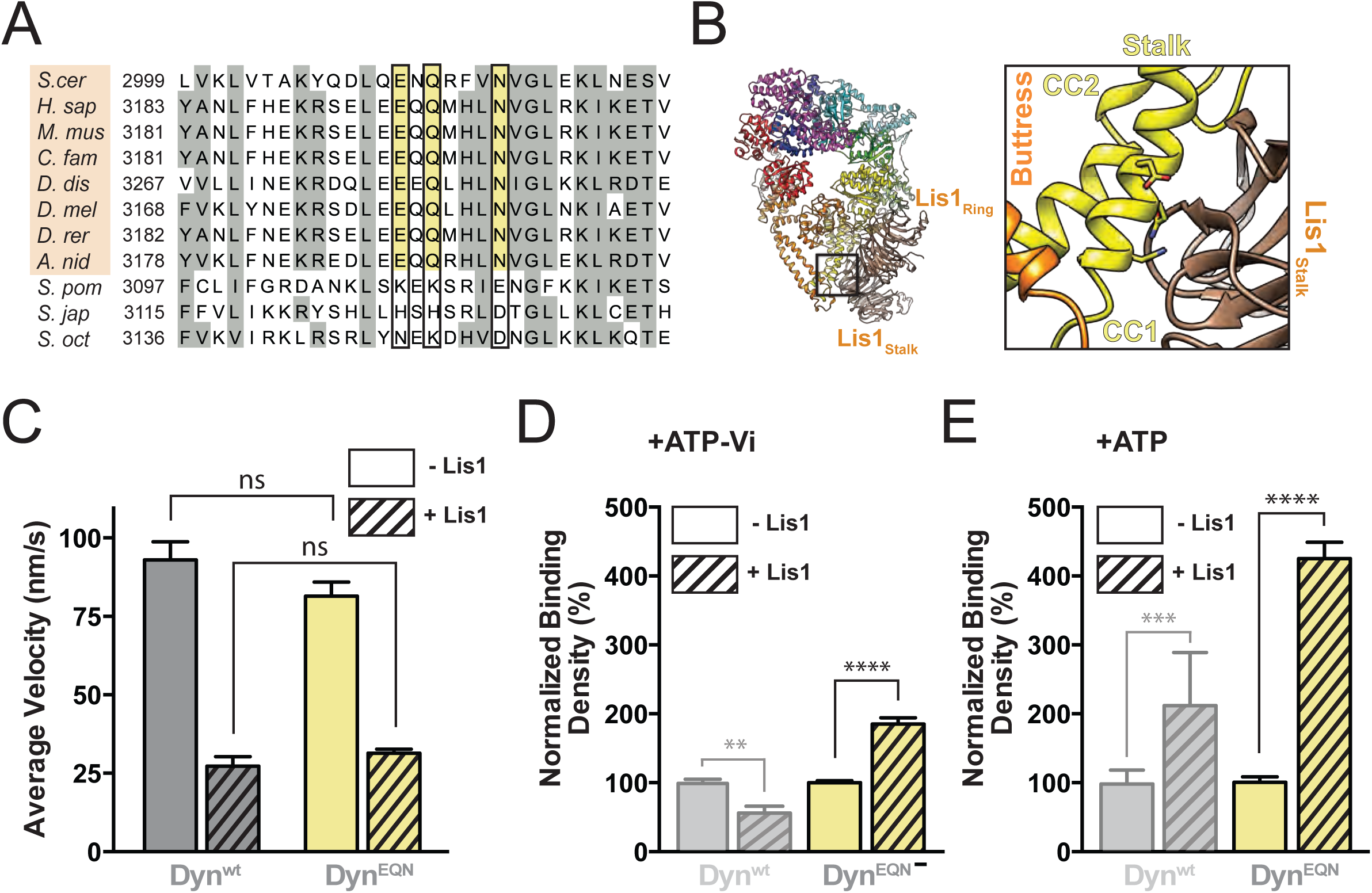
The second Lis1 binding site is required for the Lis1-induced weak microtubule binding state of dynein. (A) Sequence conservation around the putative Site_Stalk_. The figure shows the region of dynein’s stalk, extracted from a full sequence alignment of the dynein heavy chain, identified in the Dyn^WB-M^:Lis1 structure as the second binding site for Lis1. Residues with 25% conservation or higher are shaded grey. The three residues we mutated—E, Q, N, shaded in yellow—are conserved in organisms that have Lis1 orthologs in their genomes (*S. cer, Saccharomyces cerevisiae; H. sap, Homo sapiens; M. mus, Mus musculus; C. fam, Canis familiaris; D. dis, Dictyostelium discoideum; D. mel, Drosophila melanogaster; D. rer, Danio rerio; A. nid, Aspergillus nidulans*). The orange shading indicates the presence of a Lis1 ortholog. The black outline highlights that these residues are not conserved in a group of fission yeasts (*S. pom, Schizosaccharomyces pombe; S. jap, Schizosaccharomyces japonicas; S. oct, Schizosaccharomyces octosporus*) that do not have a Lis1 ortholog in their genome. (B) Atomic model of the Dyn^WB^:Lis1 complex (left) and a close-up view of Site_Stalk_. The conserved E, Q and N triad is shown in stick representation and the dynein motifs in its neighborhood are labeled. We mutated the EQN to AAA. Dyn^EQN^ is a construct that carries this mutation but has a wild-type AAA3. (C) Average velocities of Dyn^wt^ (grey) and Dyn^EQN^ (yellow) in the absence (solid bars) or presence (hatched bars) of 300 nM Lis1 (n>154 per data point). (D-E) Relative binding densities of Dyn^wt^ (semitransparent grey; these data are reproduced from Figure 1 to help in the comparison) and Dyn^EQN^ (yellow) in the absence (solid bars) or presence (hatched bars) of 300 nM Lis1 (n=12 per data point) in the presence of ATP-Vi (D) or ATP (E). The binding density in the presence of Lis1 is normalized relative to the binding density in the absence of Lis1, which was set to 100%. Statistical significance was calculated using unpaired t-test with Welch’s correction for both velocity (C) and binding density (D-E). P-values: ns, not significant; **, <0.01; ***, <0.001, ****, <0.0001. Data are shown as mean and standard error of mean.

### The second Lis1 binding site is required for Lis1-mediated weak microtubule binding

Taken altogether, our data show that Lis1 induces or stabilizes a weak microtubule binding state in dynein when AAA3 carries a Walker B mutation (mimicking an ATP-bound state) and that in this complex (Dyn^WB-M^:Lis1) a second Lis1 ß-propeller interacts with dynein’s stalk (Site_Stalk_). We wanted to test the hypothesis that Site_Stalk_ is required for Lis1 to induce the weak binding state in dynein. Having identified the putative Site_Stalk_, we mutated the conserved E3012, Q3014, and N3018 residues to alanine; we will refer to this mutant, which carries a wild type AAA3, as Dyn^EQN^. Our hypothesis was that the Lis1-induced weak binding state would no longer be available in Dyn^EQN^.

We first characterized Dyn^EQN,^s velocity and ability to bind Lis1. Dyn^EQN^ moved with the same velocity as Dyn^wt^ in a single-molecule motility assay in the absence of Lis1, suggesting that the Site_Stalk_ mutations do not grossly impair dynein’s structure or function (Figure 5C). Dyn^EQN^ binds to Lis1 with a reduced affinity compared to Dyn^wt^ (Figure S5), which is compatible with the loss of one of the two Lis1 binding sites. In addition, the velocities of Dyn^EQN^ and Dyn^wt^ were lowered by similar amounts in the presence of 300 nM Lis1 (Figure 5C), suggesting that mutation of Site_Stalk_ does not impair Lis1’s interaction with dynein at Site_Ring_.

To test our hypothesis that Site_Stalk_ is required for the Lis1-induced weak microtubule binding state we needed to measure microtubule affinity for Dyn^EQN^ under conditions that would normally lead to this state. Our prediction was that Dyn^EQN^ would remain bound to microtubules under those conditions. We showed above that, in contrast to ATP, ATP-Vi caused Dyn^wt^ to have a weaker microtubule affinity in the presence of Lis1 (Figure 1E). We interpreted this result as reflecting the similarity between Dyn^wt^ with AAA3 containing ADP-Vi, and the Dyn^WB^ mutant with AAA3 containing ATP. Therefore, we used the TIRF-based microtubule-binding assay introduced earlier (Figure 1D-G) to measure the effect of Lis1 on the microtubule binding affinity of Dyn^EQN^ in the presence of ATP-Vi. As predicted, while Lis1 had led to a 44% decrease in the affinity of Dyn^wt^ for microtubules in the presence of ATP-Vi (Figure 1E and 5D), it increased the affinity of Dyn^EQN^ by 85% (Figure 5D).

We then repeated our measurements in the presence of ATP. Under these conditions, where AAA3 can sample different nucleotide states, both weak and tight binding states should be available to Dyn^wt^ in the presence of Lis1. In contrast, the weak binding state would still be unavailable to Dyn^EQN^ due to the absence of a functional Site_Stalk_. Dyn^wt^’s affinity for microtubules in the presence of Lis1 should reflect a mixture of weak and tight binding states, while that of Dyn^EQN^ should come from tight binding states only. As a result, we would expect Dyn^EQN^’s affinity for microtubules to be higher than that of Dyn^wt^ in the presence of Lis1 and ATP. This proved to be the case; in the presence of ATP, Lis1 increased Dyn^EQN^’s binding density on microtubules two-fold further than it did for Dyn^wt^ (Figure 1D and Figure 5E).

### The second binding site for Lis1 is required for dynein localization

In yeast, dynein functions in mitotic spindle positioning (Markus and Lee, 2011a; Moore et al., 2009) Dynein does so by “pulling” on spindle pole body (SPB)-anchored microtubules from a position on the cell cortex (Figure 6A) (Adames and Cooper, 2000). To reach the cell cortex, dynein must first localize to microtubule plus ends, either via kinesin-dependent transport or recruitment from the cytosol (Carvalho et al., 2004; Caudron et al., 2008; Markus et al., 2009). Dynein’s plus-end-localization, kinesin-dependent transport, and later “off-loading” to the cell cortex all require Lis1 (Lee et al., 2003; Li et al., 2005; Markus and Lee, 2011b; Markus et al., 2009; 2011; Sheeman et al., 2003). The requirement of Lis1 for plus-end-localization and off-loading to the cortex is compatible with the prevailing model of Lis1 as a factor that promotes a tight microtubule-binding state in dynein. On the other hand, this model is at odds with Lis1’s requirement in the kinesin-dependent transport of dynein to microtubule plus ends, where the two motors are engaged in a tug-of-war that kinesin wins (Markus and Lee, 2011b; Roberts et al., 2014).

**Figure 6.**
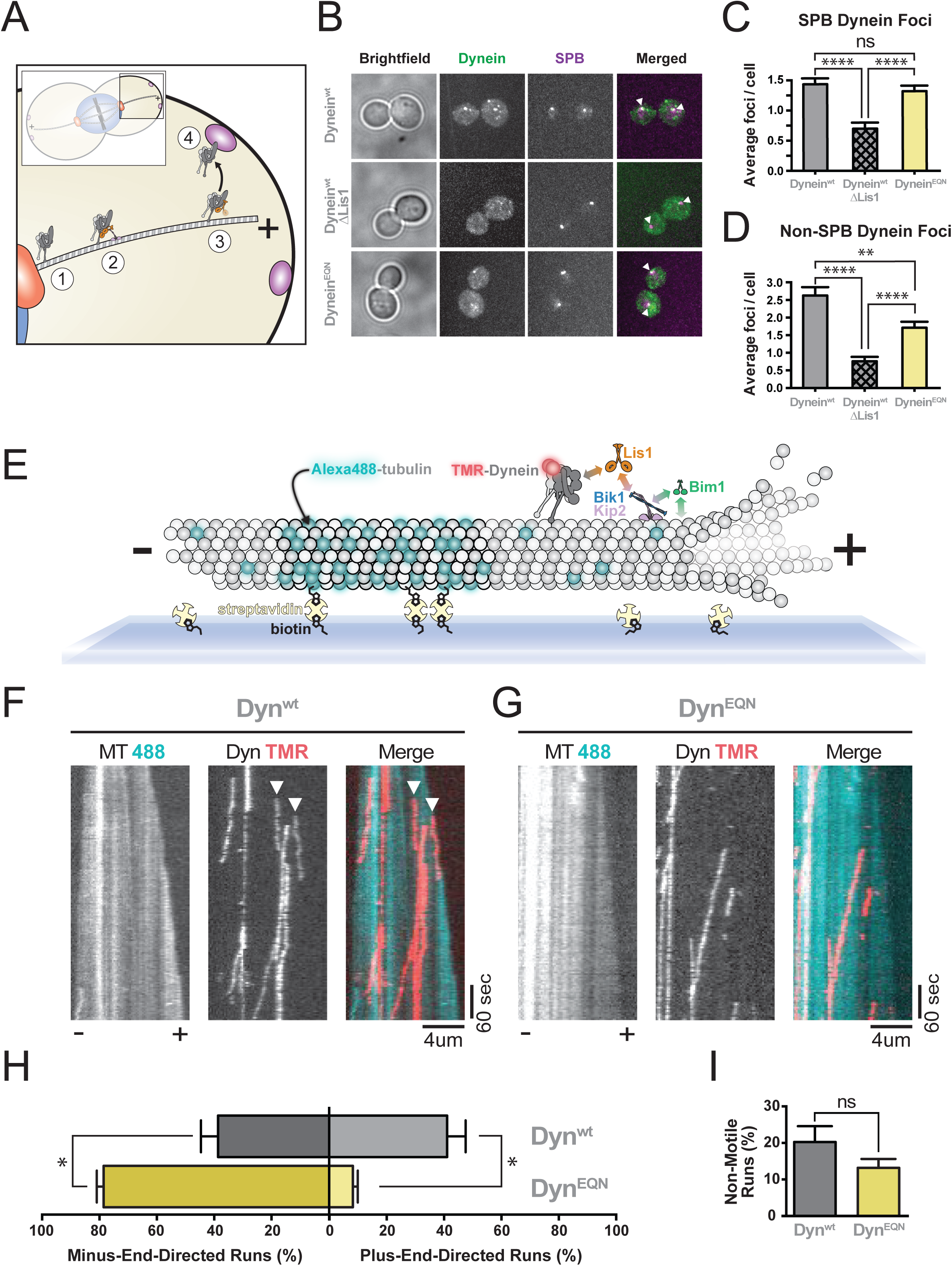
The second binding site for Lis1 is required for dynein localization in vivo and for its microtubule plus-end-directed transport by kinesin in vitro. (A) Schematic of dynein and Lis1 function in spindle positioning in *S. cerevisiae*. Dynein is first localized to the spindle pole body (1). Dynein is then transported to the microtubule plus end by a kinesin in a process that also requires Lis1 (2). Dynein is maintained at microtubule plus ends by Lis1 (3) and later “off-loaded” to the cell cortex (4), where it is activated to pull on spindle pole body-attached microtubules. This pulling helps position the mitotic spindle. (B) Dynein localization in dividing *S. cerevisiae*. First column: Representative brightfield images. Second column: maximum projections of 3xGFP-labeled dynein (Dynein^wt^ or Dynein^EQN^). Third column: maximum projections of tdTomato-labeled SPC110, a spindle pole body (SPB) marker. Fourth column: merged 3xGFP-dynein and tdTomato-SPB images. White arrowheads point to co-localized dynein and SPB signals. The strains imaged were: Dynein^wt^ (top row); Dynein^wt^ in a ΔLis1 background (middle row); Dynein^EQN^ (bottom row). Both dynein-3xGFP and SPC110-tdTomato are expressed under their endogenous promoters. (C, D) Quantification of the data presented in (B). (C) Average number of dynein foci per cell colocalized with SPBs; and (D) Average number of dynein foci per cell not colocalized with SPBs for Dynein^wt^ (grey), Dynein^wt^/ΔLis1 (hatched grey) and Dynein^EQN^ (yellow) strains (n>22 per data point). (E) Schematic representation of our in vitro reconstitution of kinesin-mediated dynein transport to the microtubule plus end. Brightly-labeled, GMPCPP-stabilized microtubule seeds are attached to the coverslip via biotinstreptavidin interactions. A dimly-labeled microtubule extension grows faster at the plus end of the seed, allowing for the microtubule polarity to be established. Addition of Dynein (grey) labeled with TMR (red sphere), Lis1 (orange), CLIP170/Bik1 (blue), kinesin/Kip2 (purple) and EB1/Bim1 (green) results in the reconstitution of plus-enddirected transport of Dynein by kinesin. Known interactions among Dynein, Lis1, Bim1, Kip2, Bim1 and tubulin are shown with double-headed arrows color-coded according to the proteins involved. The plus and minus ends of the microtubule are labeled. (F, G) Representative kymographs from the assay outlined in (E), with microtubule (MT 488) and dynein (Dyn TMR) channels shown in black and white, and the merged image in pseudocolor, for Dyn^wt^ (F) and Dyn^EQN^ (G). Plus (+) and minus (-) indicate microtubule polarity. White arrowheads point to the start of plus-end-directed runs. (H) Quantification of the percentage of plus- and minus-end-directed runs for Dyn^wt^ (Grey) and Dyn^EQN^ (Yellow). (I) Quantification of the percentage of non-motile runs. Statistical significance was calculated using Mann-Whitney test for both average number of foci per cell (C, D) and percentage of runs (H, I). P-values: ns, not significant; *, < 0.05; **, <0.01; ****, <0.0001. Data are shown as mean and standard error of mean.

To determine the cellular role of the second Lis1 binding site we identified (Site_Stalk_), we monitored dynein’s localization in vivo. To do this, we fused 3X-GFP to full-length dynein or a version carrying the EQN Site_Stalk_ mutation (dynein^EQN^) at the endogenous locus and monitored their localization. We also determined dynein localization in a Lis1 deletion (ΔLis1) background. As previously reported (Lee et al., 2003), dynein localizes to both the SPB and the cortex/microtubule plus end (Figure 6B). We could not distinguish between the cortex and microtubule plus ends in our experiments, but both sites represent localization that requires Lis1. On average, each dividing cell contains ~1.4 dynein foci at the SPB (Figure 6C) and ~2.6 dynein foci at the microtubule plus end and/or cell cortex (Figure 6D). Deletion of Lis1 leads to mislocalization of dynein as previously reported (Lee et al., 2003; Sheeman et al., 2003) (Figure 6B), with ~2- and ~3-fold reductions in dynein foci at the SPB (Figure 6C) and plus-end/cortex (Figure 6D), respectively. Interestingly, while yeast carrying the dynein^EQN^ mutant contain the same number of dynein foci at the SPB as wild type yeast (~1.3 SPB foci per cell) (Figure 6B, C), they show significantly reduced dynein at the microtubule plus end/cortex (~1.7 non-SPB dynein foci per cell) (Figure 6A, C). These data suggest that mutating Site_Stalk_ (dynein^EQN^) affects dynein’s ability to localize to the microtubule plus end and/or cell cortex. Thus, the new Lis1 binding site we identified on dynein is required for dynein’s function in vivo.

### The second binding site for Lis1 is required for dynein to be transported by kinesin

The data presented above suggest that mutating Site_Stalk_ impairs Lis1’s role in localizing dynein to the microtubule plus end/cortex. Kinesin, a microtubule-based motor, is required for the plus-end-localization of dynein in yeast (Carvalho et al., 2004; Caudron et al., 2008; Markus et al., 2009), filamentous fungi (Zhang et al., 2010), and mammalian neurons (Twelvetrees et al., 2016). Previously, we reconstituted yeast dynein plus-end-localization in vitro, showing that dynein is transported by kinesin (Kip2) to microtubule plus ends (Roberts et al., 2014). This transport requires Lis1 and two microtubule plus tip proteins CLIP170/Bik1 and EB1/Bim1 (Roberts et al., 2014). During transport, dynein and kinesin engage in a tug-of-war that results in dynein being slowly pulled towards the plus end of microtubules by kinesin, with CLIP170/Bik1 and EB1/Bim1 allowing kinesin to “win” the tug-of-war by enhancing kinesin’s processivity (Roberts et al., 2014).

We hypothesized that the weak microtubule binding state of dynein induced by Lis1 bound to Site_Stalk_ might be required for kinesin to properly localize dynein to microtubule plus ends. A prediction from this hypothesis is that the in vivo localization defect of dynein ^EQN^ (Figure 6B) could be due to kinesin’s inability to win the tug-of-war against a dynein that cannot be kept in a weak microtubule binding state. To test this, we used the assay we developed to reconstitute dynein trafficking to the plus ends of dynamic microtubules (Roberts et al., 2014) (Figure 6E). We monitored the directionality of dynein movement (i.e. towards the plus or minus ends of microtubules) in the presence of kinesin/Kip2, Lis1, CLIP170/Bik1, and EB1/Bim1 to determine whether dynein or kinesin won the tug-of-war. Dyn^wt^ was transported to the plus end in 41% of observed events (Figure 6F, H) with an average velocity of 7.1 nm/sec (Figure S6). In contrast, only 8.2% of events were plus-end-directed with Dyn^EQN^ (Figure 6G, H), but the average velocity of those events (7.7 nm/sec) was not statistically different from that seen with Dyn^wt^ (Figure S6). These results show that the second Lis1 binding site we identified on dynein is required for efficient plus-end-directed transport of dynein by kinesin to microtubule plus ends.

## Discussion

Here we report the surprising discovery that Lis1 has two opposing modes of regulating dynein, being capable of inducing both low and high affinity for the microtubule (Figure 7). This is a major revision of the current model for how Lis1 regulates dynein and has important implications for the biological roles of this ubiquitous and highly conserved dynein regulator.

**Figure 7.**
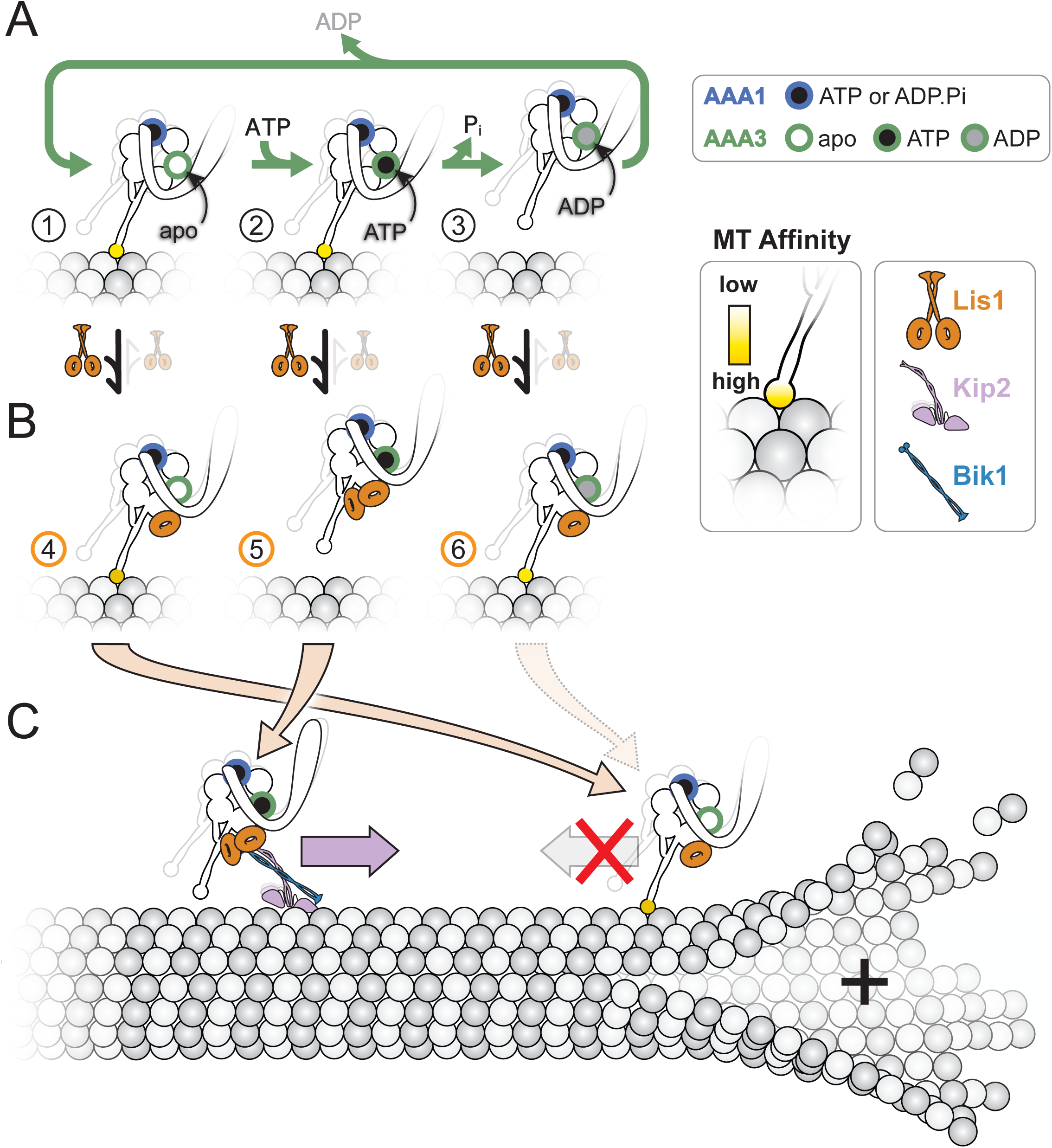
Model for the opposing modes of regulation of dynein by Lis1) (A, B) The nucleotide state of AAA3 determines which regulatory mode is used by Lis1. These panels illustrate how the ATP hydrolysis cycle at AAA3 affects the affinity of dynein for microtubules (A) and how Lis1 acts on these different states (B). We show the dynein cartoons in this figure with ATP/ADP.Pi bound at their AAA1 sites to reflect that our data suggest this is the state where Lis1 regulation is apparent. On its own, dynein has high affinity for microtubules when its AAA3 is either empty (“apo”) (A, state 1), or bound to ATP (A, state 2). Hydrolysis of ATP at AAA3 leads to the AAA3:ADP-bound, low-affinity state of the motor (A, state 3), which is expected to be the predominant state when dynein is walking along microtubules. Lis1 acts in opposite ways on states 1/3 versus 2: its ß-propeller binds in a 1:1 stoichiometry (1 Lis1 ß-propeller:1 dynein monomer) when dynein’s AAA3 is either empty or contains ADP, leading to tight binding to microtubules (B, states 4 and 6); while it binds in a 2:1 stoichiometry (2 Lis1 ß-propellers:1 dynein monomer) when dynein’s AAA3 has ATP, leading to weak binding to microtubules (B, state 5). (C) Proposed biological roles of the Lis1-mediated weak and tight microtubulebinding states of dynein. In *S. cerevisiae*, dynein is transported towards the plus-end of microtubules by the kinesin/Kip2 and CLIP170/Bik1, a process that requires Lis1. Lis1 binds to the AAA3:ATP state of dynein with a 2:1 stoichiometry (2 Lis1 ß-propellers:1 dynein monomer) (B, state 5), which keeps the motor in a weak affinity state and allows Kip2 to win the tug-of-war. Once at the plus-end of the microtubule, dynein cycles to a AAA3:ADP or AAA3:apo state (by a process not currently understood); Lis1 binding with a 1:1 stoichiometry (1 Lis1 ß-propeller:1 dynein monomer) (B, states 4 and 6) keeps the motor in a tightly bound to the microtubule in preparation for cargo loading.

Previous work from our labs and others (Huang et al., 2012; McKenney et al., 2010; Toropova et al., 2014; Yamada et al., 2008) suggested that Lis1 works by uncoupling the cycles of ATP hydrolysis at AAA1 in dynein’s motor domain from those of track binding and release at the microtubule-binding domain. This allows Lis1 to keep dynein tightly attached to microtubules and thus slows its velocity. Our new data reveal that this is only one of two opposing effects Lis1 can have on dynein.

We have now shown that Lis1 can also induce a weak microtubule-binding state in dynein. Remarkably, we discovered that Lis1’s two opposing effects are driven by different stoichiometries between Lis1’s ß-propellers and dynein, and that these are controlled by the nucleotide state of dynein’s AAA3 domain. Specifically, when AAA3 is either empty or contains ADP, a single Lis1 ß-propeller binds dynein at the interface between AAA3 and AAA4, a site we had previously identified (Huang et al., 2012; Toropova et al., 2014), and stabilizes a tight microtubule-binding state of the motor (Figures 1 and 2). In contrast, the presence of ATP at AAA3 results in a weak microtubule-binding state, where two Lis1 ß-propellers interact with dynein—one ß-propeller binds dynein at the same, canonical site at the AAA3-AAA4 junction (Site_Ring_) while the second Lis1 ß-propeller binds at a novel site on the CC1 helix of dynein’s stalk (Site_Stalk_) (Figures 1, 3 and 5). We have shown that disrupting the interaction between Lis1 and dynein at the novel Site_Stalk_ position affects the regulation of dynein by Lis1 both in vivo and in vitro. In vivo, mutating Site_Stalk_ perturbs dynein’s cellular localization (Figure 6). In vitro, preventing the binding of the second Lis1 ß-propeller eliminates the low-affinity microtubule-binding state (Figure 5) and severely impairs the transport of dynein by kinesin towards the microtubule plus end (Figure 6).

### Structural basis for the opposing modes of dynein regulation by Lis1

Our Cryo-EM structure of the Dyn^wt-M^:Lis1 complex (Figure 2) revealed a single Lis1 ß-propeller bound at SiteRing and an open ring conformation, associated with high microtubule-affinity states of dynein (Carter et al., 2011; Kon et al., 2012; Schmidt et al., 2012). This structure, however, was obtained in the presence of ATP, which normally results in dynein’s release from microtubules in the absence of Lis1. Furthermore, the linker in the Dyn^wt-M^:Lis1 complex adopts the bent, pre-power stroke position that has been observed in structures of the low-affinity ADP-Vi-bound form of dynein (Bhabha et al., 2014; Schmidt et al., 2014).

How does Lis1 stabilize the high affinity conformation of dynein’s ring under conditions that should lead to a weak microtubule-binding state? The higher resolution of the new structure presented here revealed that in addition to the interaction with AAA3 and AAA4 we identified previously (Huang et al., 2012; Toropova et al., 2014), Lis1 also interacts with AAA5 (Figure 2D). Importantly, the transition from the high affinity, open conformation of the ring to its low affinity, closed conformation involves the rotation of AAA5 relative to a rigid module comprised of AAA2-4 (Schmidt et al., 2014). This motion leads to the buttress-mediated change in the register of the stalk, and ultimately in changes in the conformation, and affinity of the microtubule binding domain (Figure 2A) (Carter et al., 2008; Gibbons, 2005; Kon et al., 2009; Redwine et al., 2012; Schmidt et al., 2014). We speculate that Lis1 prevents this conformational change and keeps dynein in a high microtubule affinity state by clamping AAA3/4 and AAA5 together, thus blocking the ring rearrangements necessary for dynein to adopt a low microtubule affinity state. We were unable to test this hypothesis here because the region of AAA5 where Lis1 interacts with dynein had the lowest local resolution in our structure (Figure S2H, I).

The most surprising result was the structure of the Dyn^WB-M^:Lis1 complex, solved in the presence of ATP-Vi (Figure 3). This structure showed a second Lis1 ß-propeller bound to dynein’s stalk (Site_Stalk_) and a closed conformation of the ring associated with a weak microtubule-binding state. Due to the clamping of AAA3/4 and AAA5 by Lis1 at Site_Rin_g, the closed conformation we observed is not identical to that of human dynein-2 in ATP-Vi (Schmidt et al., 2014). Instead, we observed that while the ring adopts an overall closed conformation, the positions of its AAA domains are displaced relative to those seen the structure of human dynein-2 (Figure 3D-F).

How do the two Lis1 ß-propellers prevent dynein from switching to a high-affinity state upon binding to a microtubule? Normally, dynein binding to microtubules triggers a conformational change in its microtubule-binding domain that alters the register of its coiled coil stalk and ultimately results in the open, high-affinity conformation of dynein’s ring (Carter et al., 2008; Gibbons, 2005; Kon et al., 2009; Redwine et al., 2012; Schmidt et al., 2014). The sliding of the two helices in the stalk (CC1 and CC2) changes their position with respect to the ring (Schmidt et al., 2014) (Figure 2A). In terms of the Lis1 binding site on CC1 (Site_Stalk_), sliding of CC1 relative to CC2, would bring Site_Stalk_ closer to Site_Ring_. Our structural analysis suggests that the precise distance between Site_Stalk_ and Site_Ring_ may play a role in Lis1’s ability to stabilize dynein’s weak microtubule-binding state. Our Dyn^WB-M^:Lis1 structure showed that Site_Ring_ is closer to Site_Stalk_ than in Dyn^wt-M^:Lis1 (Figure 5C-E). This small difference in conformation likely allows for the interaction between the Lis1 ß-propellers bound to Site_Stalk_ and Site_Ring,_ as suggested by the continuous density we observed for the two Lis1 ß-propellers (Figure 3A,B). This interaction between the two Lis1 ß-propellers could not take place in Dyn^wt-M^:Lis1 because the two Lis1 binding sites in dynein are too far apart (Figure 4F,G). We hypothesize that bridging of CC1 and the ring by the two Lis1 ß-propellers bound to dynein when AAA3 contains ATP prevents CC1 from sliding towards its high-affinity register. Future experiments will test this idea by disrupting the interaction between the two ß-propellers, but this will require a higher resolution map where the rotational orientation of the ß-propellers becomes unambiguous.

### Both modes of Lis1 regulation are required for dynein function

In *S. cerevisiae* and other organisms, some of Lis1’s known functions have been difficult to reconcile with its reported molecular function to induce a tight microtubule binding state in dynein. Below we discuss how our new model for Lis1 regulation of dynein can resolve these apparent contradictions.

As described above, dynein’s function in spindle positioning during mitosis in *S. cerevisiae* is composed of multiple steps. These include (1) SPB localization, (2) transport to the microtubule plus end, (3) off-loading to the cell cortex, and (4) pulling on SPB-attached microtubules to position the spindle (Markus and Lee, 2011a; Moore et al., 2009) (Figure 6A). By imaging dynein localization in vivo and reconstituting plus-end-directed transport in vitro we were able to probe the role of Lis1’s new dynein binding site (Site_Stalk_) and its function in promoting weak microtubule binding. Our data showed that less dynein was found at microtubule plus ends/the cortex in dynein with a mutant Site_Stalk_ (Figure 6B,D). We reasoned that this was likely due to a defect in dynein transport to microtubule plus ends, which requires a complex that includes dynein, kinesin/Kip2, and Lis1 (Roberts et al., 2014). Indeed, when we reconstituted plus-end-directed transport in vitro with Site_Stalk_ on dynein mutated, we found that less dynein moved in the plus end direction (Figure 6F-H). While keeping dynein in a weak microtubule affinity state would be beneficial for plus-end-directed transport, other dynein functions in yeast require high affinity microtubule binding, such as being retained at microtubule plus ends (Markus et al., 2011; Sheeman et al., 2003) (Lee et al., 2003). In this context we propose that Lis1 engages dynein only at Site_ring_.

The dual functionality of Lis1 we have uncovered also provides a possible explanation for the cellular function of Lis1 in other organisms, some of which have been previously difficult to reconcile with a model where Lis1 exclusively promotes high-affinity microtubule binding by dynein. For example, the fact that decreased levels of Lis1 slow the velocity of acidic organelles in mouse neurons (Pandey and Smith, 2011) and mRNAs in *Drosophila* embryos (Dix et al., 2013) points to a role for Lis1 in reducing dynein’s affinity for microtubules (and thus increasing dynein velocity). We propose that in these cases both Lis1 ß-propellers bind to dynein. A number of other studies have shown that altering Lis1 levels leads to reduced cargo transport (Klinman and Holzbaur, 2015; Moughamian et al., 2013; Shao et al., 2013; Smith et al., 2000; Yi et al., 2011), perhaps also due to the new mode of Lis1 regulation we describe here. However, other functions for dynein are more aligned with Lis1 increasing dynein’s affinity for microtubules. These include stabilizing dynein at microtubule plus ends (Li et al., 2005; Splinter et al., 2012), facilitating cargo loading onto dynein at microtubule plus ends (Egan et al., 2012; Lenz et al., 2006; Moughamian et al., 2013), aiding in the dynein-mediated trafficking of high-load cargo (McKenney et al., 2010; Reddy et al., 2016; Yi et al., 2011), and inhibiting or slowing motility of some dynein cargo (Vagnoni et al., 2016). We propose that in these cases Lis1 is bound to dynein by a single ß-propeller at Site_Ring_.

### How are the two modes of Lis1 activity regulated?

A central question for the future is to determine how Lis1 switches between these two modes of regulating dynein. The underlying basis for this switch must be regulating the nucleotide state at AAA3. There are several possible mechanisms we envision that could achieve this. Local fluctuations in the concentration of ATP or ADP could alter the rate of hydrolysis at AAA3, potentially making Lis1 regulation tunable to different cellular locations or events. Other dynein-binding partners, including Lis1 itself, could also play a role in controlling ATP turnover at AAA3. For example, NudE/NudEL binds both dynein and Lis1 and has been implicated in many Lis1-dependent functions (Cianfrocco et al., 2015). Finally, an intriguing possibility is that a backward force exerted on dynein (such as that coming from a pulled cargo) could influence the nucleotide state of AAA3 (Nicholas et al., 2015). If a backward force on dynein promoted an apo state in AAA3, this would favor the binding of a single Lis1 ß-propeller (at dynein’s Site_Ring_), and would maintain dynein in a high microtubule affinity state, which would be advantageous when dynein is working against a high load cargo. Future work will be required to understand this complex process.

## Author Contributions

MED, MAC, ZMH, SLRP and AEL designed the experiments; MED, MAC, ZMH, and PTT performed the experiments; MED, MAC, ZMH, SLRP and AEL interpreted the data; and MED, MAC, ZMH, SLRP and AEL wrote the manuscript.

## Acknowledgements

We thank Elizabeth Kellog (UC Berkeley) for the distance comparison script, the Cryo-EM facility at UCSD, Arshad Desai for the use of his labs spinning disk confocal microscope, and the Physics computing facility at UCSD, and Wei-Lih Lee (U Mass Amherst) for yeast strains. This work used the Extreme Science and Engineering Discovery Environment (XSEDE) for computing allocations (MCB160079 to ZMT and MCB140257 to AEL), which is supported by National Science Foundation grant number ACI-1548562. MED is a Howard Hughes Medical Institute Postdoctoral Fellow of the Jane Coffin Childs Memorial Fund for Medical Research. MAC is a Howard Hughes Medical Institute Fellow of the Damon Runyon Cancer Research Foundation. ZMH is supported by a NSF graduate fellowship. SRP and AEL are supported by grant R01GM107214 from the National Institutes of Health. SRP is a Howard Hughes Medical Institute-Simons Faculty Scholar.

## Accession Numbers

Cryo-EM maps and Rosetta models for the structures of Dyn^wt^:Lis1 (PDB ID: D_100022734 1) and Dyn^WB^:Lis1 (D_1000227343) have been deposited in the wwPDB.

## Methods

### KEY RESOURCES TABLE

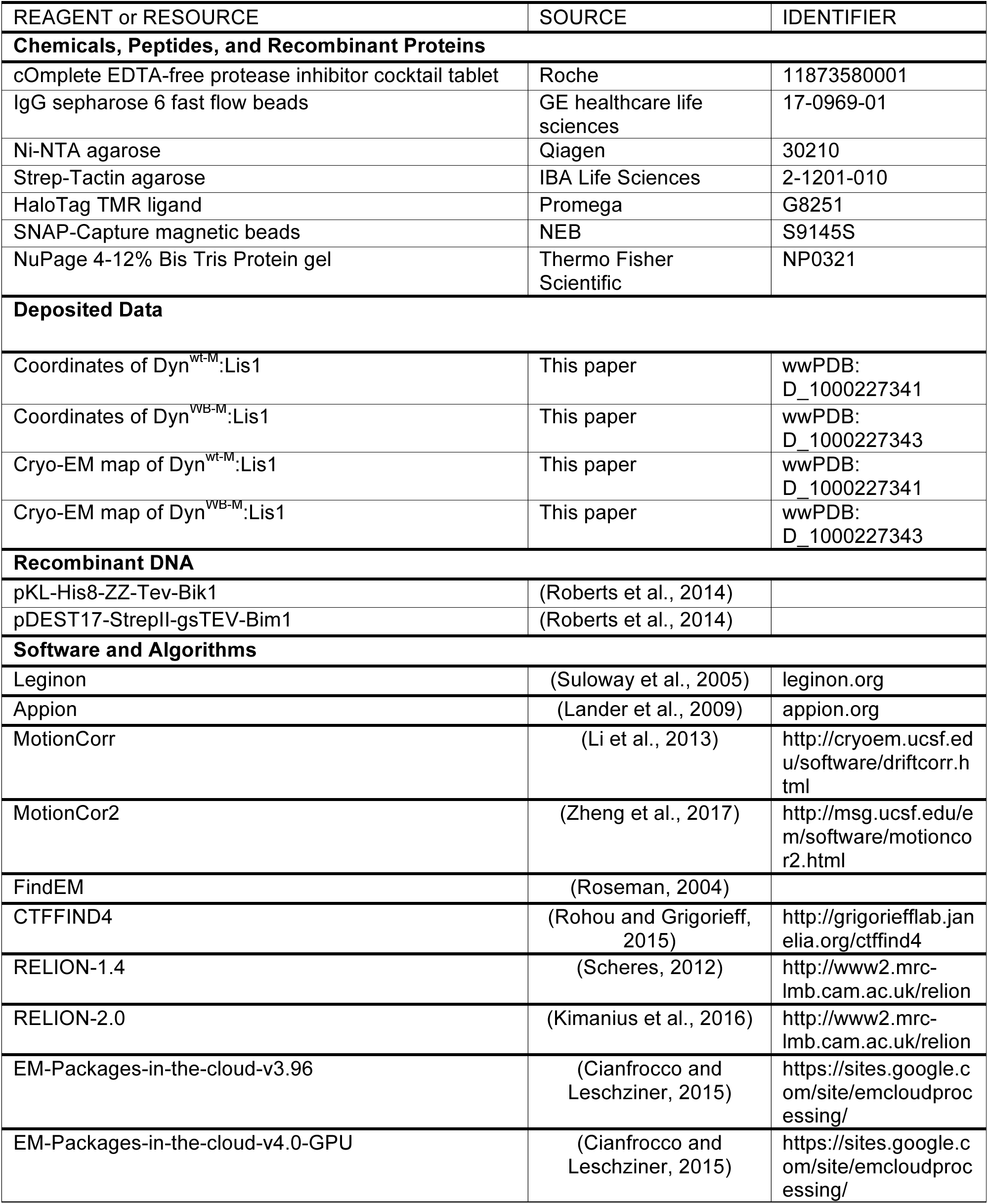

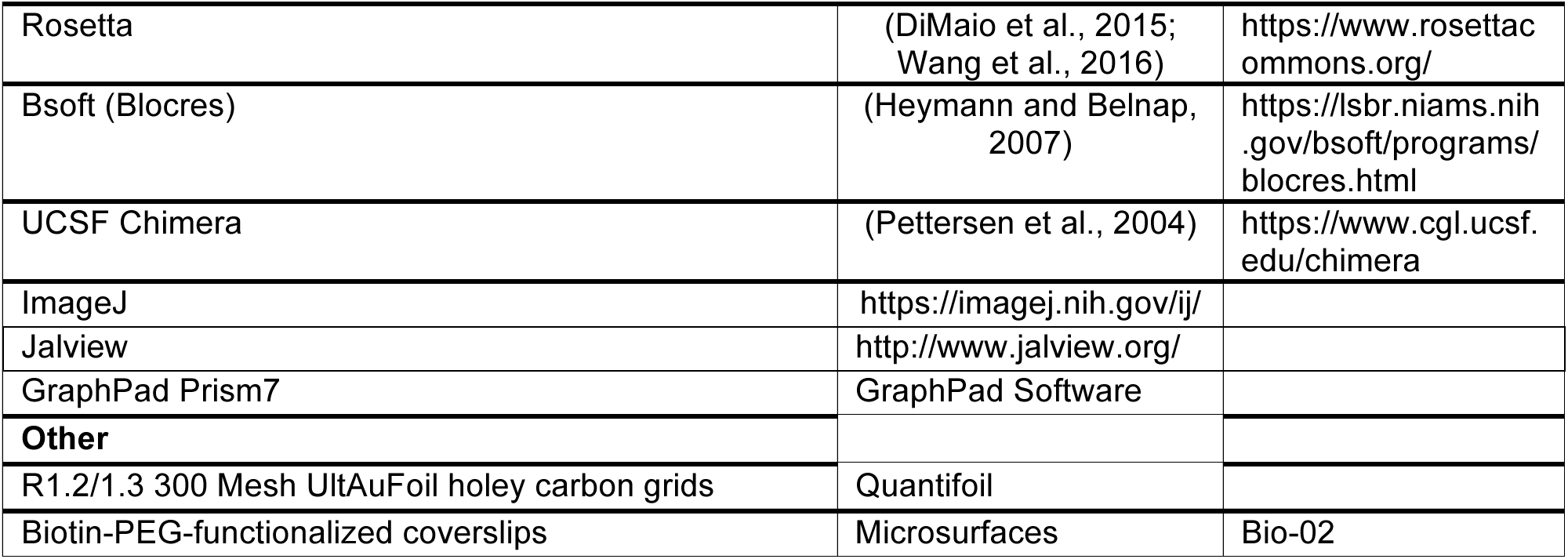

#### CONTACT FOR REAGENT AND RESOURCE SHARING

Further information and requests for reagents can be directed to, and will be fulfilled by Andres Leschziner or Samara Reck-Peterson.

#### EXPERIMENTAL MODEL AND SUBJECT DETAILS

##### Yeast strains

*S. cerevisiae* strains used in this study are listed below. The endogenous genomic copies of *DYN1, PAC1, NDL1, KIP2, TUB1* and *SPC110* were modified or deleted using PCR-based methods as described (Gietz and Woods, 2002; Longtine et al., 1998). Standard lithium acetate method (Gietz and Woods, 2002) was used to perform transformations. Cultures of *S. cerevisiae* for protein purification were grown, harvested and frozen as described (Reck-Peterson et al., 2006).

**Strain** **Genotype**
RPY1 W303a (*MATa*, *his3-11,15, ura3-1, leu2-3,112, ade2-1, trp1-1*)
RPY799 W303a *pep4Δ::HIS5, prb1Δ, P_GAL1_-8HIS-ZZ-SNAP-gs-PAC1, dyn1Δ: cgLEU2*
RPY816 W303a *pep4Δ::HIS5, prb1Δ, P_GAL1_-8HIS-ZZ-Tev-PAC1, dyn1Δ::cgLEU2, ndl1Δ::Hygro^R^*
RPY1099 W303a *pep4Δ::HIS5, prb1Δ, P_GAL1_-8HIS-ZZ-Tev-KIP2-g- FLAG-ga- SNAP−Kan ^R^*
RPY1167 W303a *pep4Δ::HIS5, prb1Δ, P_GAL1_-ZZ-TEV-GFP-3XHA-GSTDYN1(331kDa)-gsDHA-KanR, pac1Δ, ndl1Δ::cgLEU2*
RPY1302 W303a *pep4Δ::HIS5, prb1Δ, PAC11-13xMYC-TRP1, P_GAL1_-ZZ-TevDYN1(331kDa), pac1Δ:: Hygro^R^*
RPY1385 *MATa lys2-801 leu2-Δ1 his3-Δ200 trp1-Δ63 DYN1-3XGFP::TRP1, ura3-52:: CFP-TUB 1:: URA3, SPC110-tdTomato:: SpHIS5, ura3Δ::KanMX*
RPY1547 W303a *pep4Δ::HIS5, prb1Δ, PGAL1-8HIS-ZZ-Tev PAC1^R275A,R301A,R378A,W419A,K437A^, dyn1Δ::cgLEU2, ndl1Δ::Hygro^R^*
RPY1630 W303a *pep4Δ::HIS5, prb1Δ, PGAL1-ZZ-TEV-GFP-3XHA-GST-DYN1(331kDa)^K2424A^-gsDHA-KanR, pac1Δ, ndl1Δ::cgLEU2*
RPY1635 W303a *pep4Δ::HIS5, prb1Δ, PAC11-13xMYC-TRP1, PGAL1-ZZ-Tev-DYN1(331kDa)^K2424A^, pac1Δ::Hygro^R^*
RPY1653 W303a *pep4Δ::HIS5, prb1Δ, P_GAL1_-ZZ-TEV-GFP-3XHA-GST-DYN1(331kDa)^E2488Q^-gsDHA-KanR, pac1Δ, ndl1Δ::cgLEU2*
RPY1654 W303a *pep4Δ::HIS5, prb1Δ, PAC11-13xMYC-TRP1, P_GAL1_-ZZ-Te v-DYN1(331kDa)^E2488Q^, pac1Δ::Hygro^R^*
RPY1705 W303a *pep4Δ::HIS5, prb1Δ, P_GAL1_-ZZ-TEV-GFP-3XHA-GSTDYN1(331kDa)^E3012A,Q3014A,N3018A^-gsDHA-KanR, pac1Δ, ndl1Δ::cgLEU2*
RPY1708 *MATa lys2-801 leu2 -Δ1 his3 -Δ200 trp1 -Δ63 DYN1^E3012A,Q3014A,N3018A^* - *3XGFP::TRP1, ura3-52::CFP-TUB1::URA3, SPC 110-tdTomato::SpHIS5, ura3Δ::KanMX*
RPY1713 W303a *pep4Δ::HIS5, prb1Δ, PAC11-13xMYC-TRP1, PGAL1-ZZ-Tev-DYN1(331kDa)^E3012A,Q3014A,N3018A^, pac1Δ::Hygro^R^*
RPY1717 *MATa lys2-801 leu2-Δ1 his3-Δ200 trp1-Δ63 DYN1-3XGFP::TRP1, ura3-52::CFP-TUB1::URA3, SPC110-tdTomato:: SpHIS5, ura3Δ::KanMX, pac1Δ::klURA3*

P_GAL1_ indicates the galactose-induced promoter. TEV denotes a TEV protease cleavage site. DHA and SNAP denote Halo-tag (Promega) and SNAP-tag (NEB), respectively. Amino acid spacers are indicated by g (glycine), ga (glycine-alanine), and gs (glycine-serine).

#### METHODS DETAILS

##### Protein Purification

Protein purification steps were done at 4°C unless otherwise indicated. Dynein constructs were purified from *S. cerevisiae* as described previously (Reck-Peterson et al., 2006). Briefly, liquid nitrogen-frozen yeast cell pellets were lysed by grinding with a chilled coffee grinder and resuspending in dynein lysis buffer (DLB: final concentration 30 mM HEPES [pH 7.4], 50 mM potassium acetate, 2 mM magnesium acetate, 1 mM EGTA, 10% glycerol, 1 mM DTT) supplemented with 0.1 mM Mg-ATP, 0.5 mM Pefabloc, 0.05% Triton and cOmplete EDTA-free protease inhibitor cocktail tablet (Roche). The lysate was clarified by centrifuging at 264,900 x g for 1 hr or at 125,100 x g for 2 hr. The clarified supernatant was incubated with IgG sepharose beads (GE Healthcare Life Sciences) for 1.5 hr. The beads were transferred to the column, washed with DLB buffer supplemented with 250 mM potassium chloride, 0.1 mM Mg-ATP, 0.5 mM Pefabloc and 0.1% Triton, and with TEV buffer (10 mM Tris–HCl [pH 8.0], 150 mM potassium chloride, 1 mM EGTA, 10% glycerol, 1 mM DTT, 0.1 mM Mg-ATP and 0.5 mM Pefabloc). GST-dimerized dynein constructs were labeled with 5 μM Halo-TMR (Promega) in the column for 10 min at room temperature and unbound dyes were washed with TEV buffer at 4°C. Dynein was cleaved from IgG beads via incubation with 0.15 mg/mL TEV protease for 1 hr at 16°C. For dynein monomer constructs, the TEV cleavage step was done overnight at 4°C and the cleaved proteins were concentrated using 100K MWCO concentrator (EMD Millipore) to 1.5-5 mg/mL. Cleaved proteins were filtered by centrifuging with Ultrafree-MC VV filter (EMD Millipore) in a tabletop centrifuge and flash frozen in liquid nitrogen.

Lis1 and Kip2 were purified from *S. cerevisiae* as described previously (Huang et al., 2012; Roberts et al., 2014). Lysis and clarification steps were similar to dynein purification except buffer A (final concentration: 50 mM potassium phosphate [pH 8.0], 150 mM potassium acetate, 150 mM sodium chloride, 2mM magnesium acetate, 5mM β-mercaptoethanol, 10% glycerol, 0.2% Triton, 0.5 mM Pefabloc) supplemented with 10 mM imidazole (pH 8.0) and cOmplete EDTA-free protease inhibitor cocktail tablet was used as lysis buffer. The clarified supernatant was incubated with Ni-NTA agarose (Qiagen) for 1 hr. The Ni beads were transferred to the column, washed with buffer A + 20 mM imidazole (pH 8.0) and eluted with buffer A + 250 mM imidazole (pH 8.0). The eluted protein was incubated with IgG sepharose beads for 1 hr. IgG beads were transferred to the column, washed with buffer A + 20 mM imidazole (pH 8.0) and with modified TEV buffer (50 mM Tris–HCl [pH 8.0], 150 mM potassium acetate, 2 mM magnesium acetate, 1 mM EGTA, 10% glycerol, 1 mM DTT, and 0.5 mM Pefabloc). TEV cleavage was done as described for dynein purification.

Bik1 was purified from Baculovirus as described previously (Roberts et al., 2014). Cell pellets were resuspended in buffer B (final concentration: 50 mM Tris-HCl [pH 8.5], 300 mM potassium chloride, 2 mM magnesium acetate, 5 mM β-mercaptoethanol, 5% glycerol, 10 mM imidazole) supplemented with 1% NP-40 and cOmplete EDTA-free protease inhibitor cocktail tablet and lysed using a Dounce homogenizer (15 strokes with loose plunger and 10 strokes with tight plunger). The lysate was clarified by centrifuging at 183,960 x g for 30 min. The clarified supernatant was incubated with Ni-NTA agarose for 1 hr. The Ni beads were transferred to the column, washed with buffer C (20 mM Tris-HCl [pH 8.5], 500 mM potassium chloride, 5 mM β-mercaptoethanol, 20 mM imidazole), buffer D (20 mM Tris-HCl [pH 8.5], 1 M potassium chloride, 5 mM β-mercaptoethanol, 20 mM imidazole), buffer C and buffer E (20 mM Tris-HCl [pH 7.5], 200 mM potassium chloride, 5 mM β-mercaptoethanol, 10% glycerol). Bik1 was eluted with buffer E + 300 mM imidazole and flash frozen in liquid nitrogen.

Bim1 was purified from *E. Coli* as described previously (Roberts et al., 2014). Protein expression was induced in BL-21[DE3] cells (NEB) at OD 0.6 with 0.1 mM IPTG for 16 hr at 18°C. Cell pellets were resuspended in buffer B with 1 mg/mL lysozyme, incubated for 30 min on ice and lysed by sonication. The lysate was clarified by centrifuging at 154,980 x g for 30 min. The clarified supernatant was passed over Strep-Tactin agarose resin (IBA Life Sciences) three times in a column. The resin was washed with buffer D and with modified TEV buffer. TEV cleavage was done as described for the dynein purification.

##### TIRF microscopy

Imaging was performed with an inverted microscope (Nikon, Ti-E Eclipse) equipped with a 100x 1.49 N.A. oil immersion objective (Nikon, Plano Apo). The xy position of the stage was controlled by ProScan linear motor stage controller (Prior). The microscope was equipped with an MLC400B laser launch (Agilent) equipped with 405 nm, 488 nm, 561 nm and 640 nm laser lines. The excitation and emission paths were filtered using appropriate single bandpass filter cubes (Chroma). The emitted signals were detected with an electron multiplying CCD camera (Andor Technology, iXon Ultra 888). Illumination and image acquisition is controlled by NIS Elements Advanced Research software (Nikon).

##### Single-molecule motility and binding assay on taxol-stabilized microtubules

Single-molecule motility and microtubule binding assays were performed in flow chambers assembled as described previously (Case et al., 1997) using the TIRF microscopy set up described above. No. 1-1/2 coverslips (Corning) were used for the flow chamber assembly and sonicated in 100% ethanol for 10 min to reduce nonspecific binding. Taxol-stabilized microtubules with ~10% biotin-tubulin and ~10% Alexa488-tubulin were attached to the flow chamber via biotin-BSA and streptavidin as described previously (Huang et al., 2012). Dynein was labeled with Halo-TMR for visualization. For each frame, Alexa488-tubulin and TMR-dynein were exposed for 100 ms with the 488 nm laser and 561 nm laser, respectively.

For motility assays, 1-25 pM dynein was incubated with 300 nM Lis1 or modified TEV buffer (to buffer match for experiments without Lis1) for 10 min on ice, and flowed into the flow chamber pre-assembled with taxol-stabilized microtubules. The final imaging buffer contained DLB supplemented with 50 mM potassium acetate (hence a total of 100 mM potassium acetate), 20 μM taxol, 1 mM Mg-ATP, 1 mg/mL casein, 71.5 mM β-mercaptoethanol and an oxygen scavenger system (0.4% glucose, 45 μg/ml glucose catalase, and 1.15 mg/ml glucose oxidase). Microtubules were imaged first by taking a snapshot. Dyneins were imaged every 1 sec (for wild-type dynein) or 2 sec (for mutant dyneins) for 10 min. Microtubules were imaged again by taking a snapshot to check for stage drift. Movies showing significant drift were not analyzed. Each sample was imaged no longer than 30 min.

For the single-molecule microtubule binding assay, 0.5-10 pM dynein was incubated with 300nM Lis1 or modified TEV buffer (to buffer match for experiments without Lis1) for 10 min on ice. The final imaging buffer contained DLB supplemented with 20 μM taxol, 1 mg/mL casein, 71.5 mM β-mercaptoethanol, an oxygen scavenger system, and 1 mM nucleotides (Mg-ATP, Mg-ATP/NaVO4, Mg-ADP) or 2.5 units/mL apyrase for no nucleotide condition. For high-salt experiments, 150mM potassium chloride was supplemented in the final imaging buffer. Nucleotides or apyrase were added to the incubated dynein samples immediately before flowing into the flow chamber. Dynein was incubated for an additional 10 min in the flow chamber at room temperature to reach steady-state before imaging. For microtubule decoration assays with ATP, dynein was imaged by taking a single-frame snapshot. For microtubule binding assays with other nucleotide conditions, dynein was imaged every 1 sec for a total of 5 sec. Each sample was imaged at 4 different fields of view. For each set of nucleotide conditions, the samples with and without Lis1 were imaged in two separate flow chambers made on the same coverslip in order to minimize binding density variation due to nonspecific binding.

##### Cryo-EM sample preparation

Protein samples were thawed quickly and kept on ice prior to grid preparation. For Dyn^wt-M^:Lis1, the sample was prepared using the following steps: dynein and Lis1 were incubated on ice for 10 min, after which ATP was added to a final concentration of 5 mM and the sample was incubated on ice for another 10 min. This resulted in both dynein and Lis1 at a final concentration of 0.75 μM in buffer (50 mM Tris–HCl [pH 8.0], 150 mM potassium acetate, 1 mM EGTA, 1 mM DTT, and 5 mM Mg-ATP).

For Dyn^WB-M^:Lis1, the sample was prepared using the following method: dynein and Lis1 were incubated on ice for 10 min, after which Mg-ATP/NaVO4 was added to a final concentration of 1.2 mM and the sample was incubated on ice for another 10 min. This resulted in both dynein and Lis1 at a final concentration of 0.75 μM in buffer (50 mM Tris–HCl [pH 8.0], 150 mM potassium acetate, 1 mM EGTA, 1 mM DTT, 1.2 mM Mg-ATP/NaVO_4_).

After incubation, both samples were treated identically: 4 μl of sample was applied directly to an untreated (no glow discharge or plasma cleaning) UltrAuFoil 1.2/1.3 grid (Quantifoil) in a Vitrobot (FEI Company) kept at 100% humidity and 4°C. After applying the sample, the excess liquid was immediately blotted in the Vitrobot using a blot force of 20 and a blot time of 4 sec prior to plunge-freezing into liquid ethane.

##### Cryo-EM data collection and image analysis

Both datasets (Dyn^wt-M^:Lis1 and Dyn^WB-M^:Lis1) were collected on a Talos Arctica transmission electron microscope (FEI Company) operating at 200 keV with a K2 Summit direct electron detector (Gatan Inc.) (See also Table S2). Images were collected automatically using Leginon (Suloway et al., 2005) in super-resolution mode with a calibrated pixel size of 0.60 Å/pixel. The movies were then processed in the Appion pipeline (Lander et al., 2009) for all subsequent steps. Initial movie alignment and gain reference correction were performed with MotionCor (Li et al., 2013).

For the Dyn^wt-M^:Lis1 dataset (see also Figure S2 and Table 2), 485,102 particles were picked from 5,614 micrographs using FindEM (Roseman, 2004) with templates generated from forward projections at 25 degree angular increments of EMDB 6013 (Toropova et al., 2014) (dynein motor domain without nucleotide). Using these particle coordinates, particles were extracted from micrographs that were aligned using MotionCor2 (Zheng et al., 2017) in Relion-1.4 (Scheres, 2012) using defocus values calculated by CTFFIND4 (Rohou and Grigorieff, 2015) on MotionCor2 micrographs. Micrographs were discarded if CTF confidence fits from CTFFIND4 did not go beyond 10 Å. For initial 2D classification in Relion-1.4, particles were extracted at a box size of 80 x 80 pixels and a pixel size of 4.8 Å/pixel. These particles were classified into 200 classes over 11 iterations with a mask diameter of 210 Å From the resulting averages, 347,462 particles were selected from classes that did not have contaminating ice or gold particulates. These particles were subjected to another round of 2D classification using Relion-1.4, classifying them into 200 classes over 25 iterations with a mask diameter of 190 Å From the resulting class averages (of which a subset is shown in Figure S2B), 151,470 particles (80 x 80 pixels; 4.8 Å/pixel) were selected for 3D refinement to determine a structure at 9.96 Å using gold-standard FSC=0.143, using EMDB6016 filtered to 60 Å as an initial model. A summary of the 3D refinement and classification strategy is shown in Figure S2C. This refinement and all subsequent 3D classification and refinement routines were performed using Relion-2.0beta (Kimanius et al., 2016) on Amazon Web Services using EM-packages-in-the-cloud-v4.0-GPU (Cianfrocco and Leschziner, 2015). Next, particles were re-extracted at a pixel size of 1.2 Å/pixel (324 x 324 pixels) and subjected to 3D classification without alignment, using orientations determined in the previous 3D refinement step. After classifying into 3 groups over 15 iterations, 10 more iterations of classification were performed using a local angular search range of 10 degrees. From this classification, one class was selected for further refinement (45,219 particles) to obtain a 9.48 Å structure using gold-standard FSC=0.143 because the two other classes did not have high resolution features present. With this refined 3D structure, we classified the particles into 2 groups over 15 iterations, followed by 10 iterations of local angular search ranges of 10 degrees. This resulted in one class containing high resolution features that was used for a final round of 3D refinement (25,520 particles). After this last 3D refinement step, the overall resolution was calculated to be 7.7 Å applying a *B*-factor of -50 Å^2^ after combining the half-maps and masking during post-processing (Figure S2D). However, due to the presence of flexible regions of the structure, we calculated local resolution and filtered the map using Blocres in Bsoft (Heymann and Belnap, 2007), which displayed a range of resolutions from 6 – 10 Å (Figure S2E).

For the Dyn^WB-M^:Lis1 dataset (see also Figure S3 and Table S2), 414,277 particles were picked from 4,826 micrographs using FindEM (Roseman, 2004) with templates generated from forward projections at 25 degree angular increments of EMDB 6013 (Toropova et al., 2014) (dynein motor domain without nucleotide). Using these particle coordinates, particles were extracted from micrographs that were aligned using MotionCor2 (Zheng et al., 2017) in Relion-1.4 (Scheres, 2012) using defocus values calculated by CTFFIND4 (Rohou and Grigorieff, 2015) on MotionCor2 aligned micrographs. Micrographs were discarded if CTF confidence fits from CTFFIND4 did not go beyond 10 Å Particles were extracted using a box size of 64 x 64 pixels and a pixel size of 4.8 Å/pixel. Prior to 2D classification, all particles that had gold particulates (as defined by size and pixel values) were removed from the extracted particle stack. This produced a dataset that had 223,981 particles that were subsequently classified into 250 classes using Relion-1.4 (Figure S3B). After this classification, particles that belong to homogenous classes were selected for further 3D analysis (107,273 particles). Prior to 3D analysis, particles were re-extracted at a box size of 128 x 128 pixels and a pixel size of 2.4 Å/pixel. After performing 3D classification into 3 classes (Figure S3C) using EMDB6016 as a starting model, filtered to 60 Å, we obtained a single class that could be determined to an overall resolution 10.2 Å with a *B*-factor of -800 Å^2^ using Relion-1.4 (Figure S3D). Local resolution assessment using Blocres in Bsoft (Heymann and Belnap, 2007) displayed a range of resolutions from 9 – 13 Å (Figure S3E), however the map was filtered to a single value of 10.2 Å with a *B*-factor of -800 Å^2^ using Relion-1.4.

All figures were generated using UCSF Chimera (Pettersen et al., 2004).

##### Model building using Rosetta

The initial model for dynein was generated based on homology detection to the *S. cerevisiae dyn1* sequence using Hidden Markov Model as implemented in Hhpred (Söding et al., 2005) (https://toolkit.tuebingen.mpg.de/hhpred), using the top 3 scoring homologous models (PDB: 4RH7 (Schmidt et al., 2014), 3VKG (Kon et al., 2012), and 4AKG (Schmidt et al., 2012)). This model was split into three parts: 1) Linker -> AAA4(Stalk CC1), 2) AAA4(Stalk CC2) -> AAA5 Large, 3) AAA5S -> C-terminus, and each was refined using Rosetta with cryo-EM maps Dyn^wt-M^:Lis1 and Dyn^WB-M^:Lis1 (Wang et al., 2016) (DiMaio et al., 2015). In each case 200 models were generated, and the RMSD values for the top five models are shown in Figures S2H and S3H, with most RMSD values being <1 Å for Cα backbone atoms. Finally, the top-scoring model from each part was combined for a final refinement to ensure inter-domain contacts were satisfied.

##### Cα distance calculations

For Cα distance measurements and comparisons, atomic coordinates were used for only large and small AAA domains in the dynein ring. PDB coordinates from the previously published structure of human dynein-2 with ADP.Vi (PDB 4RH7) (Schmidt et al., 2014) were used to create a homology model using the sequence from *S. cerevisiae* with SWISS-MODEL (Biasini et al., 2014). Using these coordinates, distances between Cα atoms were calculated and displayed using UCSF Chimera (Pettersen et al., 2004). For each distance measurement the lines shown represent the distance between the Cα atoms, and the thickness of the linearly scaled with the distance between atoms. Program is available upon request to the authors.

##### Single-molecule motility assay on dynamic microtubules

Single-molecule motility assays on dynamic microtubules was performed using the TIRF microscopy set up described above. Flow-chambers were prepared as described previously (Case et al., 1997) using biotin-PEG-functionalized coverslips (Microsurfaces). Brightly-labeled, biotinylated and GMPCPP-stabilized microtubule seeds were prepared as described previously (Roberts et al., 2014). Flow chambers were incubated sequentially with the following solutions, interspersed with two washes with assay buffer (BRB80 [80 mM PIPES-KOH pH 6.8, 1 mM MgCl2, 1 mM EGTA], 0.5 mg/mL casein and 1 mM DTT): (1) 0.8% pluronic F-127 and 5 mg/mL casein in water (6 min incubation); (2) 0.5 mg/mL streptavidin in BRB80 (3 min incubation); (3) a fresh dilution of microtubule seeds in assay buffer (3 min incubation); and (4) the final imaging solution containing 2.5-5 pM dynein-TMR, 1 nM Kip2, 5 nM Bim1, 50 nM Bik1, 25 nM Lis1, 15 μM tubulin (~7.5% Alexa488 labeled and ~92.5% unlabeled), 1 mM Mg-ATP, 1 mM Mg-GTP, 0.1% methylcellulose, 71.5 mM β-mercaptoethanol and an oxygen scavenger system in assay buffer. Two-color sequential TIRF movies of microtubules and dynein were imaged every 2 sec for a total of 10 min. For each frame, Alexa488- tubulin and TMR-dynein were exposed for 100 ms with the 488 nm laser and for 200 ms with the 561 nm laser, respectively.

##### Lis1 affinity capture to determine Lis1-dynein binding affinities

Sixteen μL of magnetic SNAP-Capture beads (NEB) were incubated with increasing concentrations of SNAP-Lis1 (0-600 nM) in NuTEV buffer for 1 hour at room temperature with agitation. The supernatant was removed, the beads were washed with 1 ml of NuTEV followed by 1 ml of TEV buffer supplemented with 1 mM DTT, 0.1% NP40, 2 mM MgCl_2_, 1 mM ATP, and 1 mM VO_4_. 20 nM Dyn^(variant)-M^ was incubated with the beads conjugated to Lis1 for 30 min at room temperature with agitation. The supernatant was removed, ran on a 4-12% Bis Tris gel, and stained with Sypro Red (Thermo Fisher) to visualize the fraction of dynein depleted.

##### Sequence alignment

Protein sequences of dynein were obtained from UniProt. Sequence alignments were performed with Clustal Omega web services (McWilliam et al., 2013) and annotated using Jalview (Waterhouse et al., 2009).

##### Yeast *in vivo* dynein localization assay

To quantify dynein localization during mitosis, yeast strains containing 3xGFP-labeled dynein and tdTomato-labeled SPB marker, SPC110, were used. Overnight cultures grown from a single colony were diluted to OD_600_ of 0.1 in 10 mL YPD. Diluted cultures were then grown for 2-3 hours with rotation at 30°C to reach mid-log phase (OD_600_ 0.6- 0.8). 100-200 μL of mid-log phase culture were spun down using a table-top centrifuge, the media was discarded and the cells were resuspended in 3 μL of phosphate-saline buffer with calcium and magnesium. The resupended cells were then added to freshly-made synthetic complete media agarose pad on glass slide. No. 1-1/2 coverslips (Corning) were placed on top of the sample and sealed with nail polish. Imaging was performed with a spinning-disk confocal (Yokogawa, CSU10) inverted microscope (Nikon, Eclipse TE2000-E) equipped with a 100X 1.40 N.A. oil-immersion objective (Nikon, Plano Apo), and equipped with an electron multiplying CCD (Andor Technology, iXon DV887). The excitation and emission paths were filtered using appropriate single bandpass filter cubes (Chroma). Images were collected for 15 x 500 nm Z-sections (7.5 μm total Z stack). For each Z section, a bright field image, a 3xGFP-labeled dynein image via 488 nm laser excitation, and a tdTomato-labeled SPB image via 568 nm laser excitation were collected. Illumination and image acquisition is controlled by iQ2.6 imaging software (Andor Technology).

#### QUANTIFICATION AND STATISTICAL ANALYSIS

##### Single-molecule motility assay

Dynein velocities and directionalities were calculated from kymographs using an ImageJ macro as described previously (Roberts et al., 2014). Each pixel corresponds to 157 nm in our single-molecule assays. Only runs that lasted at least 4 frames were included in the analysis. Bright aggregates, which were less than 5% of the population, were excluded from the analysis. Statistical analyses for velocities were done using unpaired t-test with Welch’s correction in Prism7 (GraphPad). Run termination rates were calculated by fitting the cumulative distribution of run duration with a one phase exponential decay function in Prism7. Statistical comparisons of the percentage of plus-end moving events, minus-end moving events or non-motile events were performed using Mann-Whitney test in Prism7. Exact value of n and evaluation of statistical significance are described in the corresponding figure legends.

##### Single-molecule decoration assay

Dynein binding density on microtubules was calculated using ImageJ. A minimum projection of 5 movie frames of dynein was generated to minimize counting non-specific binding events. In the microtubule decoration assay with ATP, a single-frame snapshot of dynein was used due to dynein motility in the presence of ATP. Intensity profiles of dynein spots were generated over a 5-pixel wide line drawn perpendicular to the long axis of the microtubule. Intensity peaks at least 2-fold higher than the background intensity were counted as dynein spots bound to microtubules. Bright aggregates that were 5-fold brighter than neighboring intensity peaks were not counted as dynein spots. The total number of dynein spots was divided by the total microtubule length in each field of view to calculate the binding density. Normalized binding density was calculated by dividing by the average binding density of dynein without Lis1 in each nucleotide condition. Statistical significance was determined using an unpaired t-test with Welch’s correction in Prism7. The exact value of n and evaluation of statistical significance are described in the corresponding figure legends.

##### Yeast *in vivo* dynein localization assay

Only yeast cells with large buds and two SPB foci were included in the analysis. Maximum intensity projections of the GFP-Dynein and tdTomato-SPC110 channels were generated. Dynein and SPB foci were identified using the ImageJ plugin, *Find Maxima*, and a maxima cutoff of at least 1.5-fold higher than the neighboring background pixel intensity value. Dynein foci were separated into two categories: localized at SPB and not localized at SPB. The latter category includes cortical and microtubule plus-end-localized dynein. We cannot differentiate between these two populations because microtubules were not imaged (or labeled in our strains). Statistical comparisons of the average number of dynein foci per cell were done using a Mann-Whitney test in Prism7. The exact value of n and evaluation of statistical significance are described in the corresponding figure legends.

